# Metabolic commitment and nitrogen control of diazotrophy in the diazoplast-containing diatom *Epithemia adnata*

**DOI:** 10.64898/2026.04.23.720389

**Authors:** Adrian O. Sanchez, Ivana Antolin, Mauro Do Nascimento, Lara Sanchez Rizza, Alejandro S. Mechaly, Adrián López-García, Sara González-Bodí, Jaime Huerta-Cepas, Enrique Flores, Luis M. Rubio, Leonardo Curatti

## Abstract

Earth’s nitrogen cycle is central to sustaining ecosystem productivity and global biogeochemical balance. Although biological N₂-fixation is well characterized in prokaryotes and plant symbioses, in other eukaryotic lineages it remains poorly understood. Diatoms of the family Rhopalodiacea harbor diazoplasts, endosymbiotic spheroid bodies specialized for N₂-fixation. This makes these diatoms genuine N₂-fixing eukaryotes that represent a unique model for organelle evolution, parallel but distinct from haptophyte nitroplasts. Here, we report the isolation and stable cultivation of an *Epithemia adnata* strain, the sequencing of its diazoplast genome and its proteomic profile when growing diazotrophically in the light or darkness, or upon exposure to ammonium. Our analyses reveal that ammonium induced broad down-regulation of diazoplast proteins, particularly those linked to N₂-fixation, ATP synthesis, and central carbon metabolism underscoring a general regulatory commitment toward diazotrophic metabolism tightly coupled to host carbon and nitrogen status. The pentose phosphate pathway and ferredoxin–NADP⁺ oxidoreductase appear as likely source of reductant to nitrogenase. A striking enrichment of chaperones, peroxiredoxins, bacterioferritin-like proteins, and DpsA might stabilize nitrogenase and buffer against oxidative stress during light-driven diazotrophy. Importantly, we identified a plasmid-encoded GlpF as a putative glycerol transporter, pointing to glycerol-mediated host–symbiont metabolic integration in the extant symbiosis and possibly a crucial innovation during the early evolutionary stages of its establishment.

Thus, diazoplast activity is not autonomous but requires integration with host carbon and nitrogen status, establishing glycerol transport, reductant supply, stress mitigation, and nutrient-responsive regulation as pivotal mechanisms of nitrogenase activity and host integration. These findings have broad implications for biogeochemical cycling, organellogenesis, and synthetic biology strategies aimed at engineering N₂-fixation in crop plants.

**Significance:** N_2_-fixing eukaryotes are increasingly recognized as abundant algae containing bacterial-derived diazotrophic endosymbionts, representing an underappreciated component of global N cycling. Diazoplasts in rhopalodiacean diatoms represent a compelling example of such endosymbionts specialized for N_2_-fixation. By combining genomic sequencing and proteomic analysis, we demonstrate their metabolic specialization, host integration, and regulatory commitment to diazotrophy. These findings reinforce the emerging view that diazoplasts function as organelle-like entities dedicated to N₂-fixation, dependent on host-supplied carbon while contributing fixed N in return. Beyond giving evolutionary insights into organellogenesis, this work establishes a framework for translational applications, such as engineering N₂-fixation into agricultural plants. Such advances could reduce reliance on synthetic fertilizers, influence biogeochemical cycles, and promote sustainable food production.

## Introduction

Earth’s carbon and nitrogen cycles are essential for maintaining the planet’s fertility and habitability. Nitrogen, the fourth most abundant element required for life (after oxygen, carbon and hydrogen), often limits the growth and productivity of terrestrial and aquatic ecosystems, despite the vast atmospheric reservoir of N₂ gas. For their nutrition, humans, most animals, and fungi acquire N from organic sources. In contrast, plants and many bacteria rely on inorganic N compounds. A much smaller group of microorganisms, specifically certain Bacteria and Archaea known as diazotrophs, can access atmospheric N₂ directly through a specialized process called biological N_2_-fixation (BNF) (1). BNF is a fundamental component of Earth’s N biogeochemical cycle, playing a crucial role in offsetting N losses caused by the microbial reduction of inorganic N back to atmospheric N₂, primarily through processes such as canonical denitrification and anaerobic ammonium oxidation (anammox). Nitrogen-dependent CO₂ fixation, particularly in the vast oligotrophic oceans, not only sustains marine trophic networks but also exerts a significant influence on global climate dynamics (2).

It is widely accepted that all eukaryotic organisms descend from ancestral lineages in which at least two partners engaged in symbiotic interactions. Mitochondria originated from an α-proteobacterial ancestor that underwent significant genome reduction over evolutionary time, optimizing its function for efficient aerobic respiration and ATP synthesis, among other essential roles. Plastids, key organelles in photoautotrophic eukaryotes, are thought to have arisen from the fusion of a cyanobacterium-like ancestor with a phagotrophic eukaryote, enabling the development of oxygenic photosynthesis and CO₂ fixation in plants and algae. The metabolic demands of at least one partner appear to have driven the close interactions that ultimately led to organelle formation (1, 3).

In contrast, most mutualistic relationships involving BNF remain facultative, despite numerous adaptations that support mutual benefit. In plants, symbiotic interactions, particularly those between legumes and rhizobia, are of major scientific and agricultural significance and have been the focus of intensive investigation. The formation of N_2_-fixing nodules is a highly complex process, involving intricate molecular exchanges and signaling pathways between the plant and bacterial partner. Current understanding of the mechanisms underlying symbiosis suggests that the formation of these nodules resulted from a coevolutionary process. Notably, rhizobium-legume symbiosis is re-established in each plant generation through the selective recruitment of compatible soil bacteria by the host plant (4).

Whereas in plants several N_2_-fixing symbioses are well-characterized, the study of cyanobacterial symbionts and endosymbionts in protists remains comparatively underdeveloped, despite their recognition for several decades. Nonetheless, it is evident that protist symbioses encompass a broad spectrum, from transient associations with extracellular cyanobacteria to more stable relationships involving intracellular endosymbionts capable of N_2_-fixation (1, 5–6). The global and widespread distribution of diazotrophic diatom symbioses in marine environments, long overlooked, has highlighted their significance in Earth’s N biogeochemical cycle (7–8).

Diatoms of the family Rhopalodiaceae, especially those of the genera *Rhopalodia* and *Epithemia*, contain 1–4 (and up to 16) spheroid bodies (SB) per cell (9), which confer the ability to fix N_2_ (10). SBs are vertically transmitted during vegetative growth; they are not acquired through environmental infection and cannot survive outside the diatom host, indicating a stable endosymbiotic relationship (11). Additionally, SBs are uniparentally inherited during diatom sexual reproduction, a key trait in the evolutionary transition from endosymbiont to organelle and, certainly, a putative model system for studying organelle evolution (12, 13).

Recently, the rhopalodiacean SB (14) and the related UCYN-A, an endosymbiont of the alga *Braarudosphaera bigelowii* (15), have been regarded as diazoplasts and nitroplasts, respectively, and postulated as early stages of N₂-fixing organelles. A key discovery for that interpretation was the demonstration that UCYN-A/nitroplast appears to import up to 28% of its proteome (368 proteins) from *B. bigelowii*. These are proteins encoded in the host nuclear genome, for which a conserved C-terminal region might represent part of a targeted import mechanism (15). In contrast, diazoplasts in *Epithemia* spp. showed less integration: no gene transfer from the endosymbiont to the host has been detected *E. pelagica*, and only six host proteins of unknown function were found in the *E. clementina* diazoplast, suggesting distinct evolutionary pathways for nitroplast and diazoplast organellogenesis (14).

Metabolic integration is a defining feature of endosymbiosis and organellogenesis. From the earliest (meta)genome projects on nitroplasts (16) and diazoplasts (17) and further supported by more recent studies (8, 18–20), it became evident that both lack most of the genes required for photosynthesis, indicating dependence on carbon supplied by the algal host. Direct evidence of fixed C and N exchange between the host and the endosymbiont/N₂-fixing organelle has also been demonstrated (19, 21).

The aim of this study is to elucidate the metabolic regulation of diazoplasts. We report the isolation and stable cultivation of an *E. adnata* strain, the sequencing of its diazoplast genome, and proteomic analyses under light and dark conditions, as well as with N₂ or ammonium as nitrogen sources. Protein profiling revealed strong commitment to N_2_-fixation, ATP synthesis, and central C metabolism, closely aligned with *nif* gene regulation. These findings unravel the regulatory features of diazoplast metabolism, with potential relevance to broader questions in biogeochemical cycling, organelle evolution, and synthetic biology.

## Results

### Isolation and culture of *Epithemia adnata*

Diazotrophic eukaryotic algal holobionts are generally difficult to grow or non-cultivable under laboratory conditions, which has slowed research progress (10, 14, 15). In this study, we report the isolation of the *E. adnata* strain LB21 from a freshwater shallow lake in Buenos Aires, Argentina. Species identification was confirmed by morphological characters observed with light and electron microscopy (Fig. 1) (22). This diatom typically contains 4–8 diazoplasts, symmetrically positioned on either side of the nucleus (Fig. 1 *G-J*). Several monoalgal cultures were established and routinely maintained under laboratory conditions for the past five years. While most cultures initially displayed cell lengths of up to ∼60 µm, continuous subcultivation led to the expected shortening (23) to a mean value of 41 ± 7 µm. Notably, cultures remained viable for at least 15 months under cold temperature (4 °C) and dim light, and could resume growth producing a population of dividing cells predominantly of the longest observed size (Fig. 2 *A*). This represents a significant technical advance that can facilitate laboratory-culture-based research using this challenging group of organisms.

**Fig. 1.**
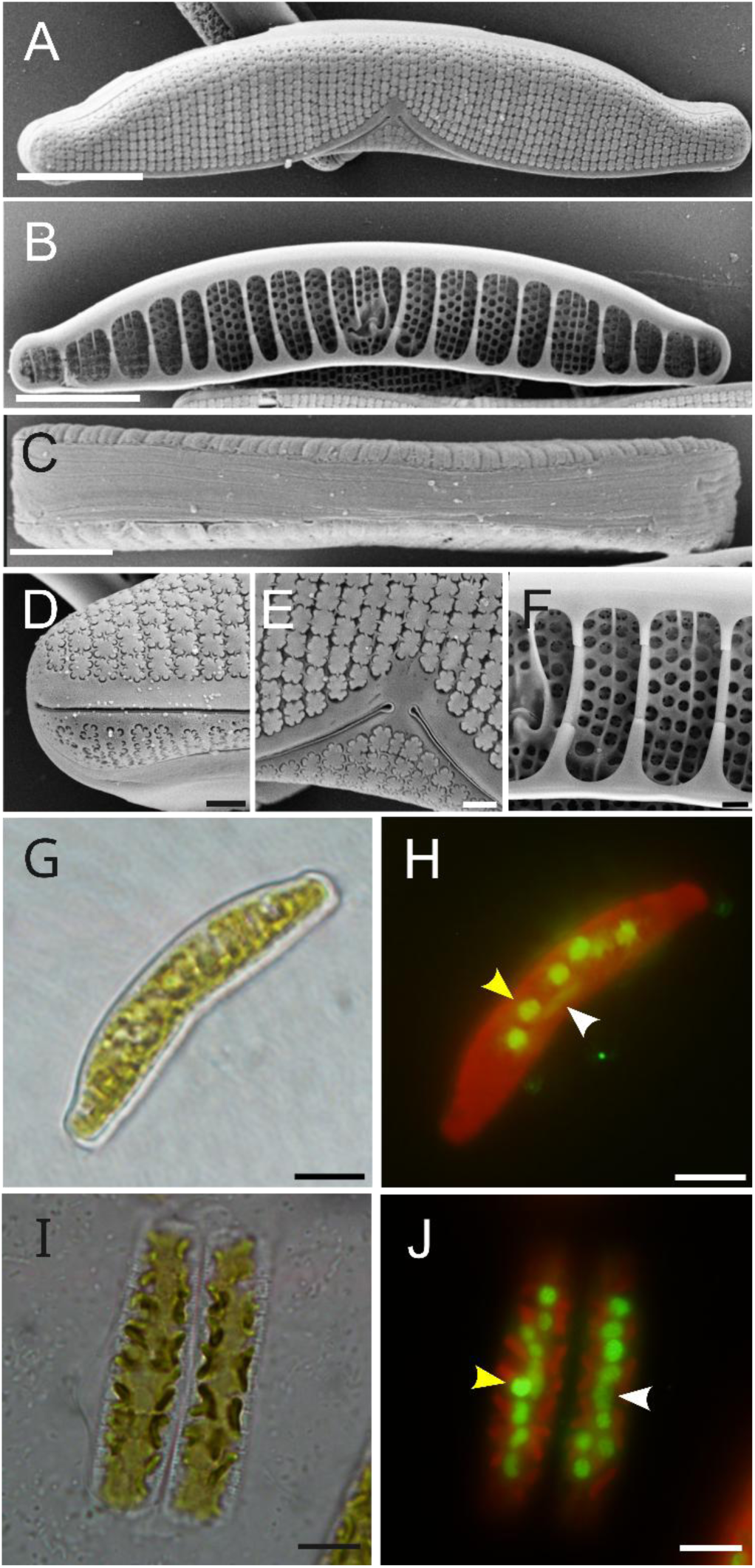
Scanning electron *(A-F)* and light *(G-J)* microphotographs of *E. adnata* LB21. *(A, B)* Whole frustules in external and internal valve views, respectively. *(C)* Girdle view. *(D, E)* External features showing the raphe ends and typical lunate areolae. *(F)* Internal features showing the valvocopula, the raphe canal and the internal wall of areolae. *(G-J)* Microphotographs in light brightfield (*G, I*) or epifluorescence (*H, J*) for valve (*G, H*) or girdle (*I, J*) views. In panels (*H, J*) DNA was stained with SYBR Green I, showing the lenticular diatom nucleus (white arrowhead) at the ventral side of the valve and diazoplasts (one indicated by a yellow arrowhead) at the dorsal side of the valve. Bars represent 10 µm in *(A-C, G-J)* and 1 µm (*D-F*).

**Fig. 2.**
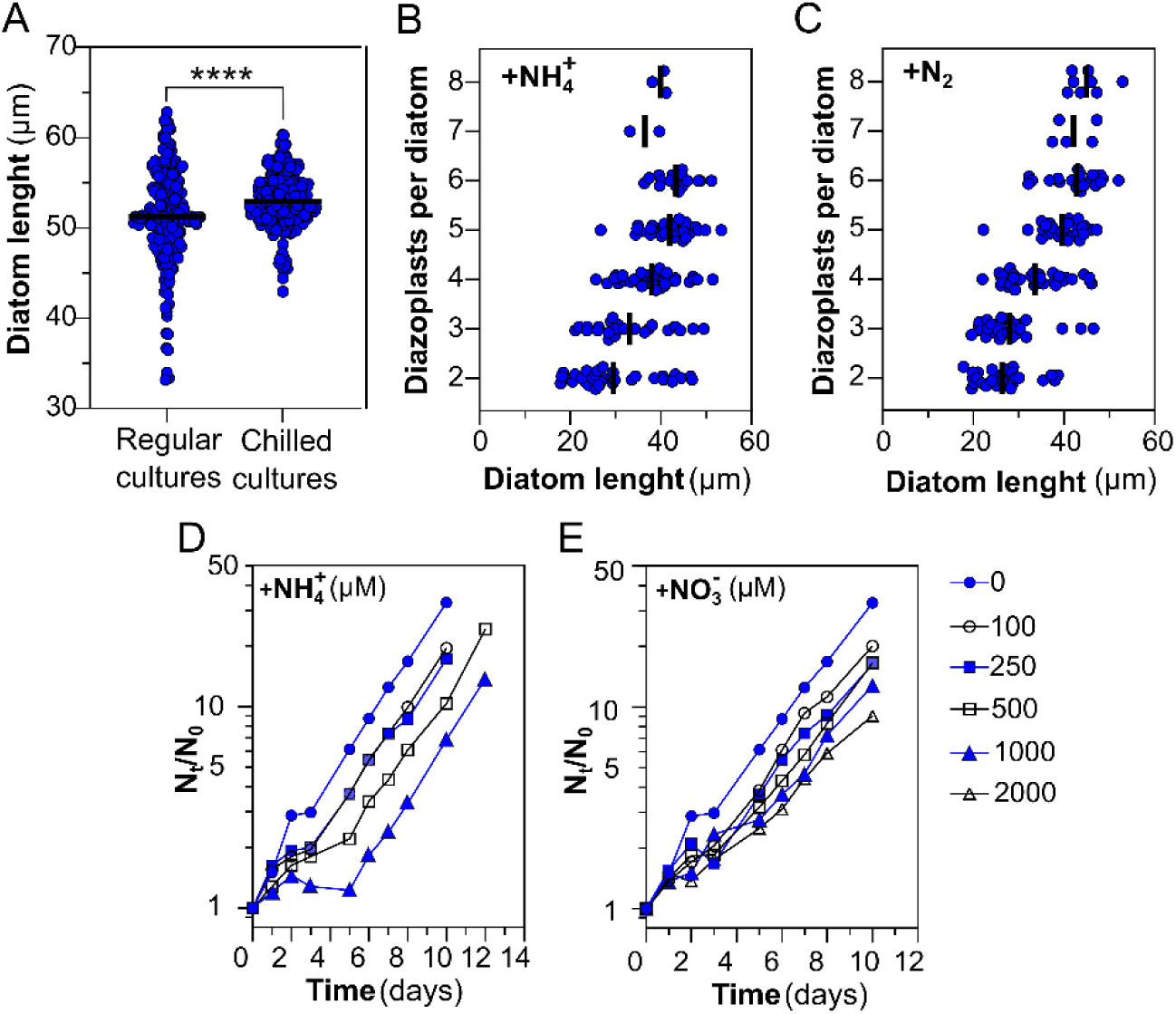
Growth characteristics of *E. adnata* LB21 in laboratory culture. (*A)* Individual diatom length based on measurements of 224 and 315 cells from regular or chilled cultures, respectively. Statistical analysis was performed using the Mann-Whitney U-test (****, *p* <0.0001). (*B, C*) diatom length in cultures supplemented with intermittent re-supply of NH_4_^+^ for 28 days (*B*) or maintained diazotrophic (*C*), determined from 151 or 153 individual cells, respectively. (*D, E*) Growth curves using NH_4_^+^ (*D*), NO_3_^-^ (*E*), or N_2_ (no ammonium or nitrate added, same data in *D* and *E*, to facilitate a comparison). Each data point represents the mean of 8 independent diazotrophic cultures (*D, E*) or 4 independent cultures supplemented with NH_4_^+^ or NO_3_^-^. Error bars in *D* and *E* were omitted for clarity.

Cultures maintained under intermittent re-supplementation of NH₄⁺ for 28 days (to avoid growth inhibition by a single larger dose, see below) exhibited a very similar number of diazoplasts per diatom cell (4.0 ± 1.4) compared to diazotrophically cultivated cells (4.3 ± 1.6) (Fig. 2 *B-C*).

### Physiology of N_2_-fixation in *Epithemia adnata*

*Epithemia adnata* showed a diazotrophic growth rate of µ = 0.32 ± 0.03 · day^-1^ (n = 8 independent cultures (Fig. 2 *D*, *E*). In the presence of NH_4_^+^, a concentration-dependent lag phase was observed of up to about 5 days at 1 mM, followed by growth with a specific growth rate similar to that observed with N_2_ (Fig. 2 *D*). In contrast, in the presence of NO_3_^-^, cells showed a slightly lower specific growth rate, around µ = 0.30 · day^-1^ (Fig. 2 *E*) at the highest concentration tested, but not a delay in the onset of the exponential growth phase of growth. Cultures fixed N₂ primarily during the daytime, as determined by acetylene reduction assays (Fig. 3 *A*). The addition of NH₄⁺ caused a short-term, transient decrease in acetylene reduction activity (Fig. 3 *B*), with recovery coinciding with NH₄⁺ depletion from the culture medium (Fig. 3 *C*). Only when combined with darkness did activity fall below the detection limit of our method (Fig. 3 *D*). In contrast, cells acclimated to NH₄⁺ exhibited a much slower increase in activity after transfer to NH₄⁺-free medium and showed a more permanent decrease in acetylene reduction upon re-exposure to similar concentrations of this N source (Fig. 3 *D*).

**Fig. 3.**
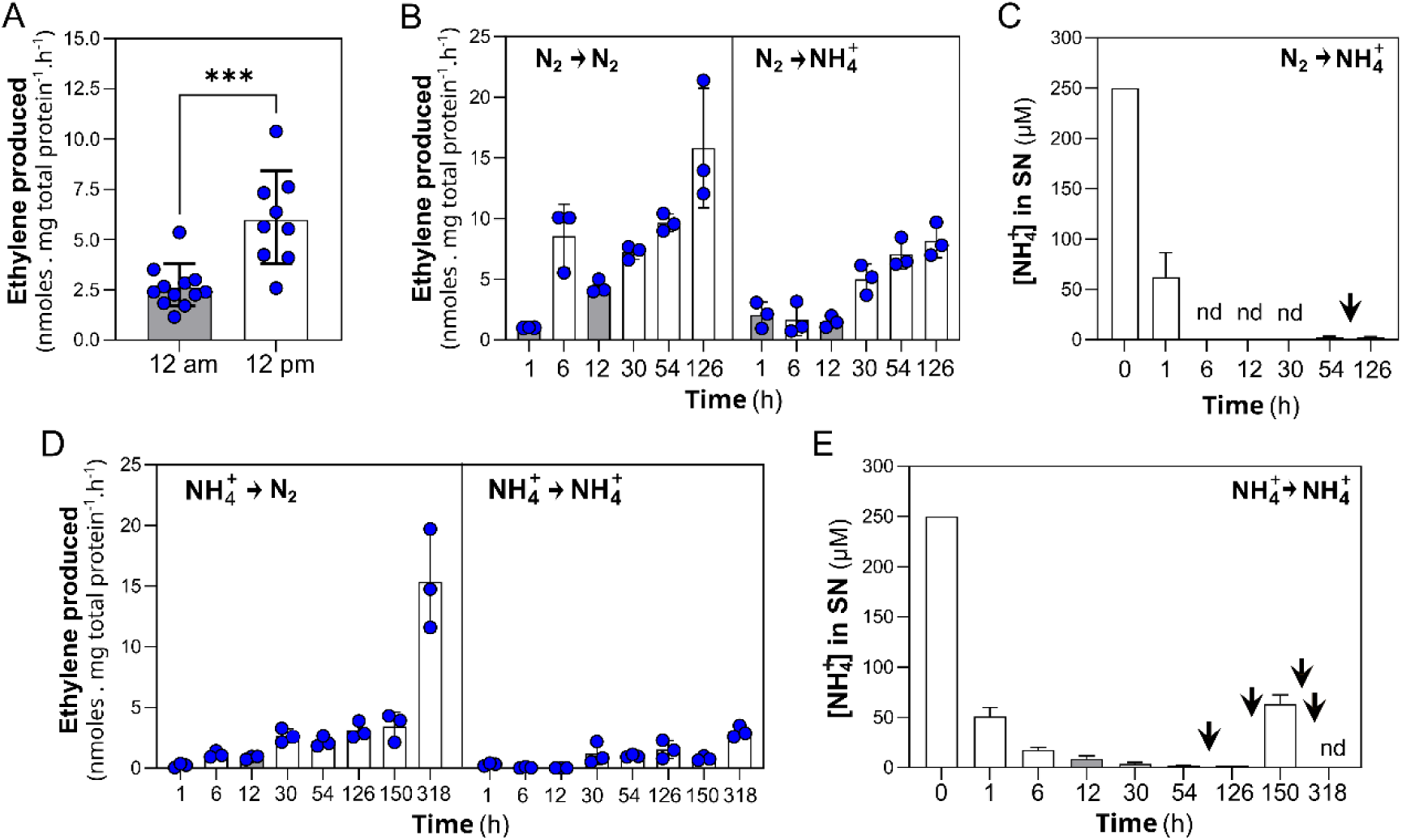
Acetylene reduction in *E. adnata* cultures. (*A*) C₂H₂ reduction assay performed on cultures sampled at the midpoint of either the dark phase (12 a.m.) or the light phase (12 p.m.) of a 12 h light/12 h dark cycle. Each data point represents an independent culture. For statistical analysis, outliers were removed using the ROUT test, followed by an unpaired t-test (***, *p* < 0.001). (*B, D*) Time course of C₂H₂ reduction assay of (*B*) cultures transferred from N_2_ to N_2_ (left side) or from N_2_ to 250 µM NH_4_^+^ (right side) in the presence of 500 µM HEPES/KOH, pH 7.5; or (*D*) cultures transferred from NH_4_^+^ to N_2_ (left side), or from NH_4_^+^ to NH_4_^+^ (right side). (*C, E*) Time course of the NH₄⁺ remaining in the culture medium at the indicated time points, showing the transitions from N₂ to NH₄⁺ (*C*) and from NH₄⁺ to NH₄⁺ (*E*). Black arrowheads in *C* and *E* mark the time of NH₄⁺ resupply at the initial concentration of 250 µM. Mean values and standard errors in panels (*B*–*E*) are based on three independent cultures. Gray shading in the bars of each panel indicates time points from the dark phase of the photoperiod. nd, not detected.

The accumulation of *E. adnata* proteins for nitrogenase structural components, NifH and NifDK (Fig. 4 *A, B*), and for the biosynthesis of its active-site FeMo-cofactor, NifB and NifEN (24) (Fig. 4 *D*) showed no apparent changes during the diel cycle (Fig. 4), suggesting that modulation of acetylene reduction activity does not occur affecting the level of these enzymes. On the other hand, statistically significant down-regulation of NifH, NifDK and NifEN was observed in diatom cultures supplemented with NH₄⁺ (Fig. 4), which could account for the observed difference in acetylene reduction activity. A detailed time course of NifH and NifDK accumulation in response to NH₄⁺ supplementation and light/dark cycles further confirm these results (*SI Appendix*, Fig. S1).

**Fig. 4.**
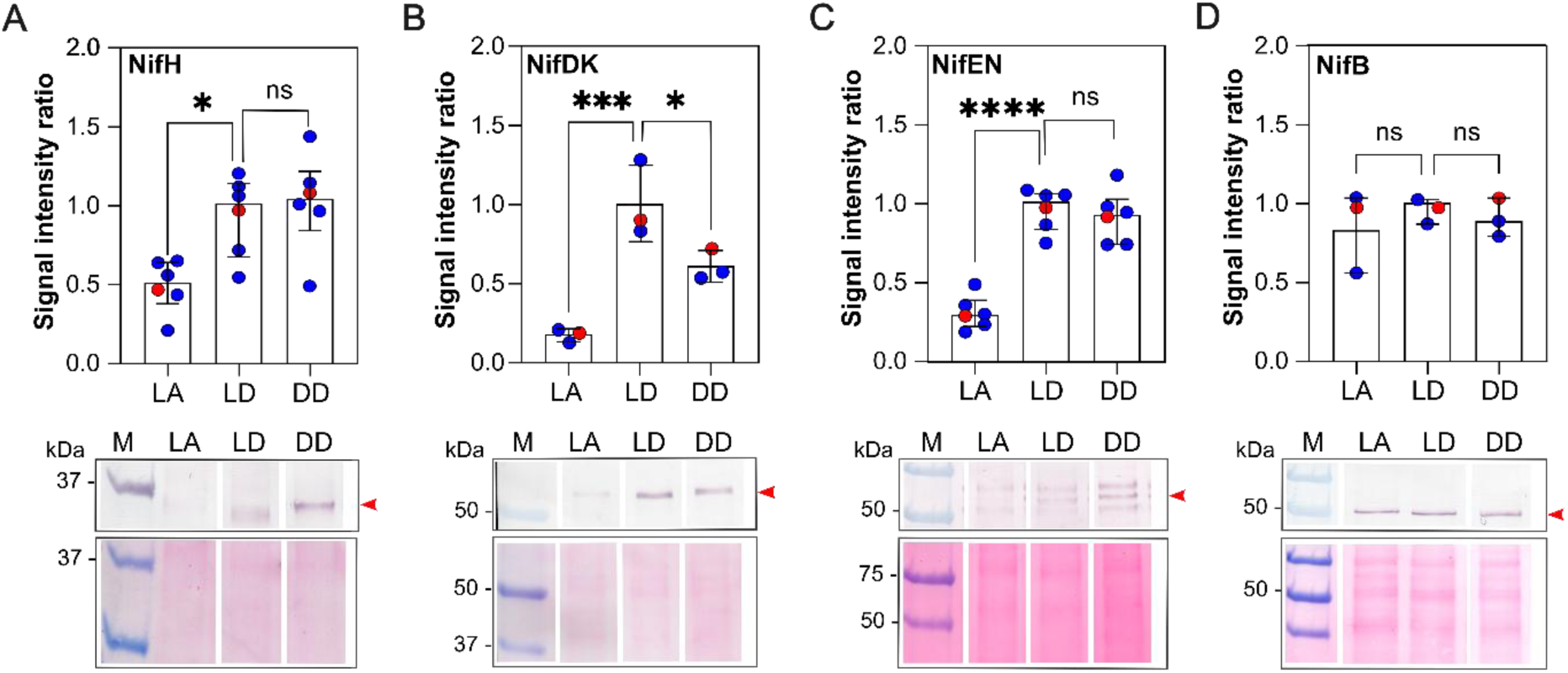
Immunoblot analysis of *E. adnata* Nif protein levels in response to light and NH₄⁺ supplementation. Reactivity tested with antibodies against the *A. vinelandii* (*A*), NifH (*B*), NifDK, (*C*) NifEN, and (*D*) NifB proteins is shown. Each lower panel displays a representative immunoassay (upper subpanel) and its corresponding protein loading control by Ponceau S staining (lower subpanel). Red arrowheads in the upper subpanel indicate the estimated specific signal for each Nif protein. Each upper panel shows the densitometric quantification of signals. Each data point corresponds to an independent assay, with the signal intensity from cultures of the reference condition (central bar labeled LD, light-diazotrophic) set to 1 for fold-change calculations. Experimental conditions were LA (left, light-NH₄⁺) and DD (right, dark-diazotrophic). Data points in red correspond to the representative blots shown in the lower panels. Statistical significance was assessed using the Kruskal–Wallis test followed by Dunn’s test (ns, not significant; *, p < 0.05; ***, p < 0.001; ****, p < 0.0001). Light/dark or N_2_/NH₄⁺ culture conditions as stated in caption to Fig. 3.

The fact that cultures maintained under intermittent re-supplementation of NH₄⁺ exhibited a number of diazoplasts per diatom cell very similar to diazotrophic cultures (Fig. 2 *B, C*) indicates that changes in C₂H₂ reduction activity and Nif protein accumulation in response to NH₄⁺ are due to regulation of gene expression or protein levels, rather than to a short-term NH₄⁺-dependent uncoupling of the diazoplast and diatom host cell cycles.

### Genome sequence of the diazoplast of *Epithemia adnata* LB21

Using a combination of Illumina and Oxford Nanopore sequencing technologies (ONT), we assembled four circular DNA molecules of 2,782,194 bp, 120,460 bp, 36,719 bp, and 5,640 bp from genomic DNA of *E. adnata*. These correspond to the diazoplast chromosome, chloroplast and mitochondrial genomes of the diatom, and a diazoplastic plasmid, respectively (*SI Appendix*, Fig. S2). Phylogenetic analysis of the diazoplast chromosome revealed that the closest relative of the *E. adnata* LB21 diazoplast currently available in public databases is from an *E. adnata* diatom isolated from the Clark Fork River, Montana, USA (20). This straińs diazoplast was recently sequenced and shares 99.95% sequence identity based on OrthoANI analysis. The next closest relative is the *E. turgida* diazoplast (17), with 97.51% sequence identity by a similar analysis (*SI Appendix*, Fig. S3). In addition, these diazoplasts, along with that of *R gibba*, harbor plasmids of 5,643 bp, 5,707 bp, and 6,464 bp, respectively. These plasmids show sequence identities of 99.26%, 87.23%, and 78.83% to the *E. adnata* LB21 plasmid sequenced in this study, as determined by BLASTn analysis.

A comparative genomic analysis of the *E. adnata* diazoplast with its closest free-living phylogenetic relative, *Crocosphaera subtropica* ATCC 51142 (NC_010546.1), showed a substantial reduction of genome size of 44%. In close agreement with the first analysis using the *E. turgida* diazoplastic genome (17), and the most recent comparison using the *E. adnata* diazoplastic genome (19Bon2 SB) (20, 25), most gene losses correspond to genes for photosynthesis (PSI and PSII; see *SI Appendix*, Fig. S4,), C fixation, HCO_3_^-^ transport, and chlorophyll biosynthesis. We also confirmed partial loss of genes for vitamins/cofactors biosynthesis (biotin, folate, pentothenate/CoA, pseudocobalamin, piridoxal 5-phosphate, and riboflavin); partial loss of genes for glycolysis and the tricarboxylic acid cycle; genes for nitrate transporters, Mo-co biosynthesis and nitrate or nitrite reductases; Fe^3+^, K^+^, Co^2+^, Ni^2+^ and Mn^2+^ and SO_4_ ^2-^ transporters; and a diminished set of genes for chemotactic response, sensing (including redlight) and signaling, and transport of some sugars and amino acids.

The *E. adnata* diazoplast genome shows a cluster of *nif* genes with the gene order *nifTZW-cysE-nifBSHDKENXW*, but it lacks *nifMUFYJ*, *nafY*, *fdxN*, and any other additional known *nif* gene. The cluster is associated with *hesAB*, *feoB1/2A*, *modB*, *fdxH*, and *fdxB*, as well as to a cluster of genes for a Ndh respiratory electron transport chain, all of which might encode N_2_ fixation-associated functions. This *nif*-gene arrangement appears to be highly conserved among the diazoplast genomes sequenced so far, as well in the *B. bigelowii* nitroplast (*SI Appendix*, Fig. S5) and some phylogenetically related free-living cyanobacteria, as also reported before (17, 25) with a few, but remarkable exceptions.

Diazoplasts and their free-living phylogenetically related likely ancestors would encode in their genomes *nifP* (*cysE*), for a serine acetyltransferase, which is a key enzyme in cysteine biosynthesis, a presumably source of S for the biosynthesis of the nitrogenase system [Fe-S] (26). No *nifP* (*cysE*) was identified in the nitroplast genomes. The diazoplasts of both *E. adnata* strains, as well as those of *E. turgida* and *R. gibba* would encode a truncated, likely inactive, copy of *nifU*, between *nifS* and *nifH*. All known diazoplast genomes would encode two sulfur utilization factor (SUF) gene clusters (27) at distinct genetic loci: *sufSDCBR* and *sufES*, suggesting *sufBCD* and or/*sufBCSDE* can either fully or partially compensate for the loss of *nifU* and/or *nifSU* for the biosynthesis of [Fe-S] clusters of the nitrogenase system. On the other hand, the genome of the free-living reference strain *C. subtropica* also encodes one copy of *nifSU* and one of *suf* genes, with a slightly different architecture (*sufRBCSDE*).

Unlike nitroplasts, diazoplasts retain a conserved arrangement of putative genes homologous to the Fe²⁺ transporter genes *feoBA*, for iron transport across the plasma membrane, located between *fdxH* and *modB*, which is also present in their likely free-living ancestor *C. subtropica* ATCC 51142. Although not unique, this conserved *feo* gene arrangement is exceptionally rare, being present in fewer than 0.6% of the available genome sequences (28). This gene arrangement features a predicted FeoB protein split into distinct N-and C-terminal regions (designated NFeoB and CFeoB, corresponding to FeoB1 and FeoB2, respectively), in contrast to the most prevalent fused-domain architecture of FeoB proteins, and is followed by a downstream copy of FeoA (*SI Appendix*, Fig. S6). All diazoplast plasmids reported to date encode a second copy of the *feoAB* genes, in a relatively uncommon configuration consisting of a *feoA*-like gene and a C*feoB*-like gene. Free-living, phylogenetically related cyanobacteria carry a similar copy of *feoAB* genes in their chromosome. However, in these cases, *feoB* is most frequently present in the canonical architecture, with fused N*FeoB* and C*FeoB* domains (*SI Appendix*, Fig. S6). Diazoplasts and nitroplasts encode a highly conserved copy of the unique porin Slr1908, which mediates iron-selective transport across the outer membrane in *Synechocystis* sp. PCC 6803 (29). This porin is widely distributed among cyanobacteria and is particularly notable in unicellular marine diazotrophs (29).

Diazoplast, nitroplast and most related cyanobacterial genomes contain fused *modBC* genes (annotated here as *modB*) for the permease and ATP-binding domains of an ABC-type transporter for Mo, while the *modA* gene for the substrate binding domain is located at a separate genomic locus.

All available diazoplast plasmids encode a putative glycerol facilitator, GlpF. The *E. adnata* diazoplast GlpF shows ∼30% identity and ∼49% similarity (biochemically similar residues) to *E. coli* GlpF, and retains amino-acid positions broadly conserved among bacterial GlpF proteins, including those demonstrated to be critical for glycerol uptake in *E. coli* (30). (*SI Appendix*, Fig. S7). Homologs are present in the chromosomes of some cyanobacteria, reaching up to ∼70% identity with the *E. adnata* diazoplast copy. In contrast, *glpF* was not found in the chromosomes of some phylogenetically related free-living taxa such as *C. subtropica* and *Rippkaea orientalis*, and is also missing in the nitroplast genomes.

For NH_4_^+^ assimilation and N (C/N) sensing and control, highly conserved copies of *glnA* (glutamine synthetase) and *glsF* (annotated as *gltB* in the diazoplast genome of *E. adnata* strain LB21), encoding a ferredoxin-dependent glutamate synthase (Fd-GOGAT), were identified in the *E. adnata* LB21 diazoplast chromosome. The predicted proteins share 82% and 72% sequence identity, respectively, with homologs from *Synechocystis* sp. PCC 6803 (31). In addition, a copy of *amt1*, encoding an ammonium/methylammonium translocator highly conserved among cyanobacteria (>75% identity), was identified in the diazoplast chromosome. The N regulatory protein PII (GlnK), the global N regulator NtcA, and the PII-interacting protein X (PipX), are highly conserved across diazoplast genomes and 90%, 79% or 50% identical to those of the free-living reference strain *Synechococcus* sp. PCC 7942 (32). Consistent with the absence of genes required for nitrate assimilation, no clear homologs of NtcB, a highly conserved LysR-type transcription factor in cyanobacteria that is essential for nitrate utilization (33), were identified in diazoplast genomes.

Unlike phylogenetically related free-living N₂-fixing cyanobacteria, E. adnata LB21 and the other available diazoplast genomes lack *cphA* and *cphB* genes for cyanophicin nitrogen storage (25, 34).

The nuclear genome assembly of *E. adnata* LB21 rendered an only partially complete genome, with 89% of universal single-copy orthologs recovered (75% single-copy, 14% duplicated), 3% fragmented, and 8% missing. This assessment (*SI Appendix*, Fig. S8), using the stramenopiles_odb10 dataset from OrthoDB v10, underscores both the substantial coverage achieved and that the remaining gaps will require additional long-read sequencing to achieve a more accurate nuclear genome sequence for this diatom.

### *Epithemia adnata* diazoplast proteome

To assess the degree of commitment to N_2_-fixation, the relative abundance of diazoplast polypeptides in diazotrophically cultivated *E. adnata* during LD condition was investigated. Among the proteins predicted from the genome sequence, most were assigned to the COG groups of “unknown” or “not assigned” functions. In contrast, the majority of the 386 proteins identified in proteomic analysis fell into defined COG categories, with only a small fraction remaining unclassified (*SI Appendix*, Fig. S9). The diazoplast proteome showed a striking enrichment in molecular chaperones, particularly GroES, GroEL1, and GroEL2, which together accounted for ∼20% of the detected proteins (Fig. 5). Nitrogenase components NifHDK represented ∼6% of the proteome, with significant levels of NifW and NifX, proteins involved in the maturation of NifDK metal clusters, also detected. Additional abundant proteins included those of the pentose phosphate pathway (PPP), ATP synthase subunits, the ferredoxin-NADPH oxidoreductase PetH, a ferritin-like domain-containing protein and DpsA, a stress-induced DNA-binding protein (Fig. 5).

**Fig. 5.**
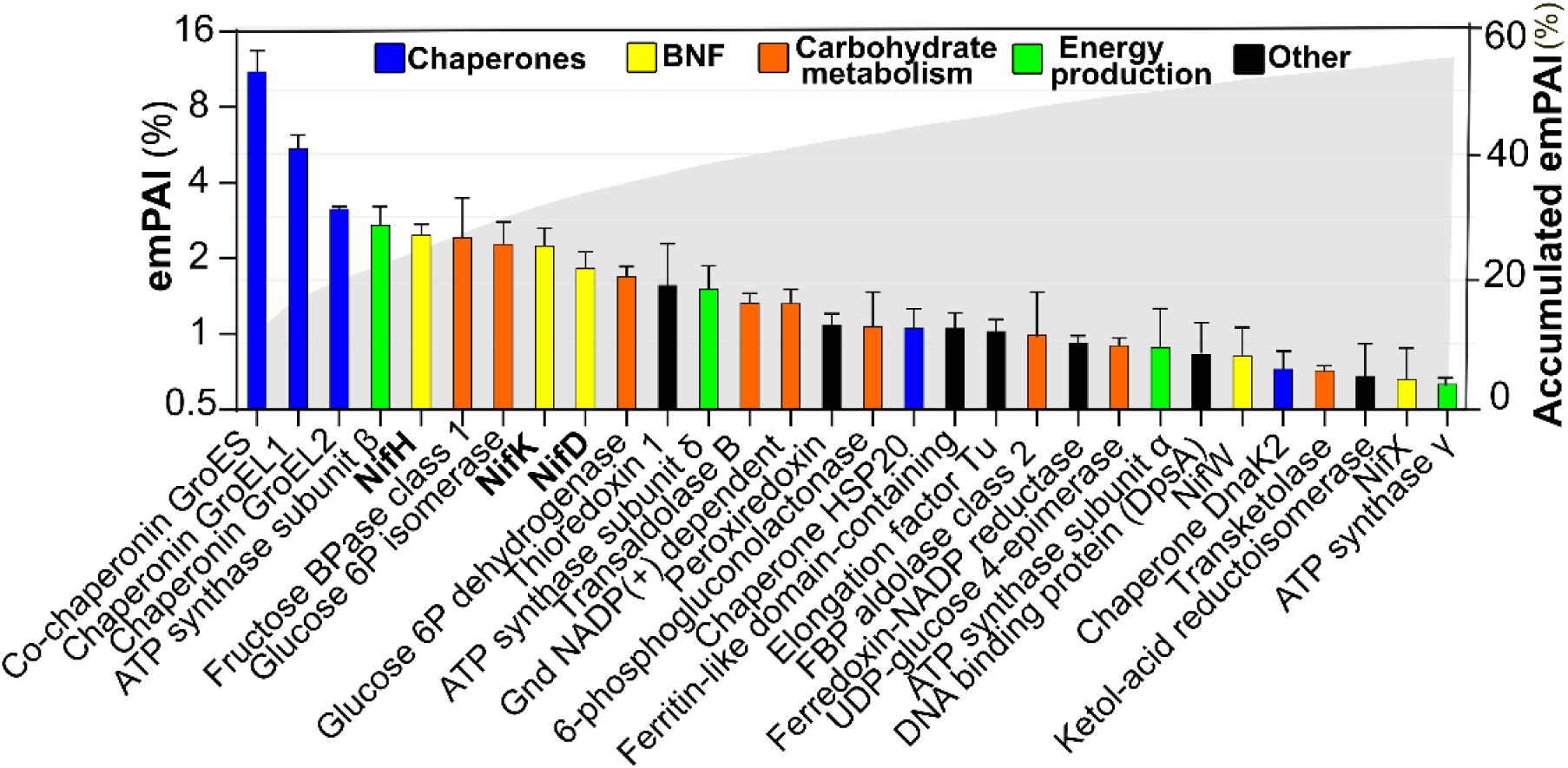
Static proteome analysis of the *E. adnata* LB21 diazoplast from diazotrophically cultivated diatoms in the light. Three independent samples were analyzed for relative abundance, calculated as emPAI (%) (35). The most abundant protein groups (molecular chaperones, nitrogenase structural proteins, carbohydrate metabolism/pentose-P pathway, and energy production/ATPase components) are color-coded. The grey shading represents the cumulative emPAI (%).

To assess regulatory mechanisms of N₂-fixation in *E. adnata*, we analyzed proteomic changes under light/dark cycling (DD condition) and in response to NH₄⁺ supplementation (LA condition). No global proteomic shift was detected in diazotrophically cultivated diatoms collected during either the illuminated or dark phases of the photoperiod. In contrast, NH_4_^+^ addition induced a substantial modification of protein expression, with a large proportion of diazoplast proteins becoming down-regulated. Despite the overall trend, most proteins exhibited only modest changes in abundance (approximately 0.5–1-fold), while a relatively small subset displayed more substantial up- or down-regulation (Fig. 6 *A*, *B*). Although proteins across most functional categories responded to NH₄⁺ supplementation, the effect was particularly pronounced in those classified under energy production and conversion, as well as in proteins of still unknown function (*SI Appendix*, Fig. S10). Notably, Nif proteins, including NifH, NifD, NifK, NifE, and NifN, showed a moderate decrease in abundance in NH₄⁺-supplemented diatoms, while remaining largely unresponsive to darkness (Fig. 6 *C*). These findings are consistent with the relative abundances previously demonstrated by immunoblot analysis (Fig. 3).

**Fig. 6.**
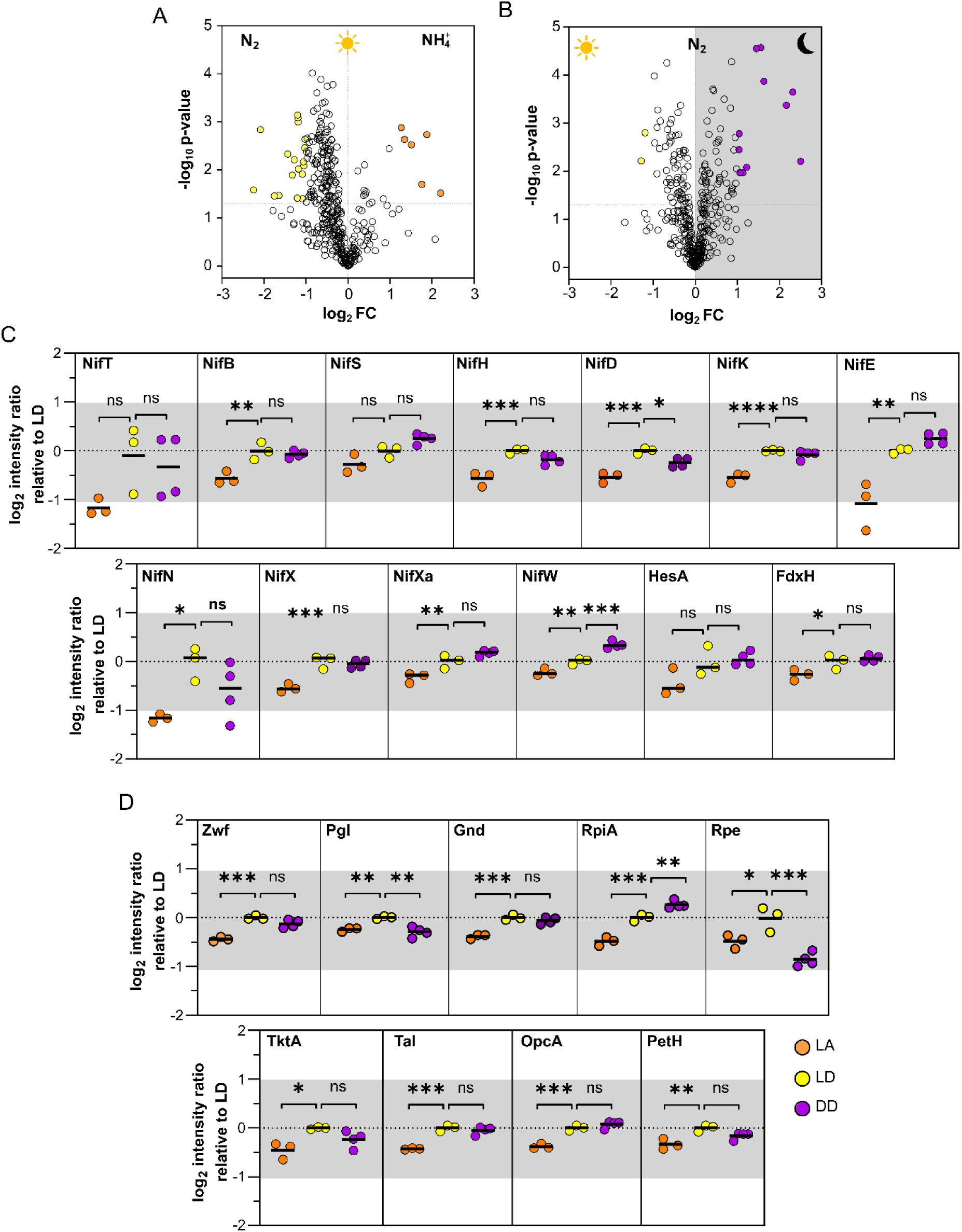
Dynamic proteome analysis of the *E. adnata* LB21 diazoplast in response to the light/dark regime and NH₄⁺ supplementation. (*A*, *B*) Volcano plots of proteins differentially expressed in (*A*) NH₄⁺-supplemented cultures in the light (LA vs. LD) and (*B*) diazotrophic cultures in the dark (DD vs. LD). Colored circles highlight the most responsive proteins whose changes in abundance are statistically supported in panels *C* and *D*. (*C*, *D*) Relative abundance of (*C*) Nif and nitrogenase accessory-proteins (known or predicted) and (*D*) pentose-P pathway proteins and PetH, which was included for comparison (see main text). Keys: LD, light-diazotrophic; LA, light-NH₄⁺; DD, dark-diazotrophic. Statistical differences of the resulting values relative to the LD condition were assessed using one-way ANOVA with Dunnett’s post hoc test (ns, not significant; *, p < 0.05; ***, p < 0.001; ****, p < 0.0001).

GlnA and GlsF, key enzymes for NH₄⁺ assimilation via the GS-GOGAT cycle, exhibited a comparable pattern: their abundances declined modestly yet significantly, to 0.75-fold (p = 0.046) and 0.78-fold (p = 0.013), respectively, while remaining insensitive to light. Proteins of the pentose phosphate pathway (PPP), together with the ferredoxin–NADPH oxidoreductase PetH, displayed expression profiles similar to those of proteins involved in N₂ fixation and NH₄⁺ assimilation under both NH₄⁺ supplementation and darkness (Fig. 6B, C), suggesting a metabolic connection. Other diazoplast-encoded proteins preferentially accumulated during diazotrophic growth, including cytochrome *f*, plastocyanin, and glucokinase, supporting NH₄⁺-dependent regulation of sugar oxidation and electron transport linked to ATP production (*SI Appendix*, Table S1). In contrast, molecular chaperones, the most abundant proteins in this dataset, appeared to be expressed largely constitutively under the tested conditions (*SI Appendix*, Fig. S11).

The circadian oscillating protein (COP23) (36) homologue EAPSB_01376 was among the most highly expressed proteins in illuminated, diazotrophically cultivated cells, showing more than a 4-fold increase compared with diatoms grown in the presence of NH₄⁺, and was also moderately up-regulated by light (*SI Appendix*, Table S1). Notably, the COP23 homologue gene contains the sequence GTA(N)₈TAC(N)₂₃TA(N)₃T located 84 nucleotides upstream of the ATG start codon (*SI Appendix*, Fig. S12). This motif represents a perfect match to the binding site of the cyanobacterial N control transcription factor NtcA in Class II promoters. While some NH₄⁺-regulated proteins, including the hypothetical protein EAPSB_00699 and PetH, also carry this sequence signature, others were associated with putative NtcA-activated Class I promoters, where NtcA may bind further upstream of the −10 determinant (not shown). In addition, several putative promoters of NH₄⁺-responsive genes exhibited degenerate binding sites (not shown), which may require NtcA co-activators for effective regulation, as previously demonstrated in certain cyanobacteria (37).

## Discussion

Although rhopalodiacean diatoms containing diazoplasts have been known for decades (7), only recently have relatively stable cultures been reported under standardized laboratory conditions (8, 12, 17, 18). Such stability is essential for establishing research models to advance knowledge of their physiology, genetics, and biotechnological potential. Diatom life cycles are typically linked to progressive cell size reduction during vegetative division, as each daughter inherits half of the frustule and forms a smaller hypotheca until sexual reproduction restores maximal size (21). Nonetheless variability exists: some species show no significant size change, suggesting either a prolonged cycle or alternative strategies that bypass sex (26). Our observations demonstrate that *E. adnata* can be maintained for at least five years under continuous subcultivation, despite some frustule shortening. Notably, chilled cultures tend to retain their apparent maximum frustule length, indicating that viability may be extended beyond its usual limit by this simple preservation adjustment. This represents a valuable technical contribution, though its applicability to other species or strains remains to be established. Abresch and colleagues (2024) likewise reported a stable culture of *E. adnata* isolated from the Clark Fork River, Montana, USA (18). Together, these complementary findings highlight the feasibility of developing laboratory models for diazotrophic diatoms, enabling multi-laboratory approaches to their physiology, an advance that has been rare in this field until now.

Diatom diazoplast genomes reported to date show a high degree of sequence identity (*SI Appendix*, Fig. S3). Especially the near-identical OrthoANI values (99.95%) observed between North and South American *E. adnata* strains underscore their minimal genomic divergence. This may reflect a deceleration in evolutionary rate following divergence from free-living relatives or, alternatively, a relatively recent dispersal between the continents via migratory waterbirds. Such avian vectors are known to connect distant habitats along flyways, with dispersal distances regularly exceeding hundreds and occasionally thousands of kilometers (38). To resolve the origin of this striking genomic similarity and evolutionary details of the group, further research, including a comparative analysis of nuclear genomes, is needed.

Comparative genomics of diazoplast-containing diatoms reveals a striking pattern of genome reduction accompanied by the selective retention of core metabolic functions. Diazoplasts consistently conserve the machinery for N_2_ fixation and central energy metabolism, particularly the pentose phosphate pathway (PPP) and ATP synthesis. This streamlined architecture underscores the evolutionary priority of maintaining reductant and energy supply for nitrogenase activity. In line with this interpretation, key genes for likely nitrogenase associated biosynthesis of [Fe-S] clusters were maintained in the diazoplast genomes, reinforcing commitment to N_2_ fixation. Namely, a copy of *nifP* for a serine acetyltransferase, which is a key enzyme in cysteine biosynthesis, a presumed source of S for the biosynthesis of nitrogenase system [Fe-S] clusters; conservation of a full set of *suf* genes, which might compensate for the loss of *nifU* in some diazoplast genomes (39), and two copies of *feoAB* genes for Fe^2+^ acquisition. Interestingly, one of the copies of *feoAB* is encoded in a small plasmid, which has been mostly overlooked in previous analyses (25).

This plasmid also encodes the glycerol uptake facilitator GlpF, which is highly conserved among diazoplast plasmids. This supports Elhai’s (2026) proposal of a central role for glycerol in metabolic integration between diatom hosts and diazoplast-containing endosymbionts (25). Once imported, glycerol would be converted into dihydroxyacetone phosphate (DHAP) by glycerol kinase and FAD-dependent glycerol-3-phosphate dehydrogenase, channeling C into glycolysis and the pentose phosphate pathway (PPP). Glycolytic intermediates are also substrates for the non-phosphorylating glyceraldehyde-3-phosphate dehydrogenase GapN, additionally producing NADPH. Glycerol-driven PPP flux, together with GapN, strengthens NADPH generation, highlighting the endosymbionts’ specialized adaptation to sustain reducing power for biosynthesis and nitrogen fixation. The proposal is further supported by the established ability of marine diatoms to synthesize glycerol, particularly in response to fluctuating salinity. Interestingly, osmolyte production in diatoms is influenced by nitrogen availability: under nitrogen sufficiency, amino acids predominate, whereas nitrogen limitation favors the synthesis of polyols such as sugars and glycerol. These metabolic and regulatory capabilities may have shaped the evolutionary origin of the associative interaction between the ancestors of Rhopalodiacean diatoms and N_2_-fixing diazoplast, likely within a marine environment (25, 40, 41). A central role of glycerol in metabolic integration is further illustrated in endosymbiotic associations, where fungi (*Rhizopus* sp.) act as suppliers and bacteria (*Mycetohabitans* sp.) as consumers (42).

Physiological, proteomic, and regulatory analyses from this study on *E. adnata* LB21 provide additional evidence for metabolic integration in the diazoplast symbiosis. The LD proteome was heavily enriched in the structural components of nitrogenase (NifHDK), proteins of the PPP, subunits of ATP synthase, ferredoxin-NADPH oxidoreductase (PetH), and bacterioferritins. The abundance of these proteins suggests a strong metabolic commitment of the diazoplast toward N_2_ fixation and related pathways for supplying reductant and ATP. Based on knowledge from cyanobacteria (43–44), electron donation to nitrogenase in diazoplasts via the PPP-PetH-ferredoxin axis appears plausible. In particular, ferredoxins such as FdxH and/or FdxB are likely candidates for electron donors, as their putative genes are interspersed among *nif* genes within the diazoplastic genome’s *nif* gene cluster.

Similar to other diazoplasts and nitroplasts, *E. adnata* LB21 fixes N_2_ predominantly during the light period, in contrast to its closest free-living cyanobacterial relatives, which typically fix N_2_ at night to protect the O_2_-sensitive nitrogenase enzyme from inactivation. This is noteworthy because, although diazoplasts have lost the O₂-evolving photosystem II of their cyanobacterial ancestors, active O₂ evolution by the host chloroplast may still interfere with N₂ fixation, raising the possibility of specialized metabolic adaptations to mitigate this conflict. Proteomic profiling under LD conditions revealed strong enrichment in molecular chaperones, thio- and peroxiredoxins, a ferritin-like domain-containing protein (bacterioferritin) and DpsA, adaptations that likely support sustained N₂ fixation under light and help buffer against constraints imposed by the endosymbiotic lifestyle. Previous studies have shown that the SUF system can substitute the *nifUS* system either for *nif*-specific or general [Fe-S] biosynthesis, especially under oxidative stress conditions (27). In this context, decreased reliance of some diazoplasts on NifU may represent an adaptive strategy, enabling N_2_ fixation in environments where oxidative stress is prevalent. Thio- and peroxiredoxins, not only could be part of an active strategy to cope with general oxidative stress (45), as it has been shown that a thioredoxin (TrxM) specifically binds the N-terminal catalytic domain of NifU in a diazotrophic cyanobacterium, suggesting that TrxM is involved in the Fe–S cluster biogenesis as well (46).

High bacterioferritin and DpsA accumulation is notable given their dual roles: storing iron for [Fe-S] and heme biosynthesis, and detoxifying excess iron to protect against O₂ and reactive oxygen species. In anaerobic organisms, or at anaerobic reaction centers, bacterioferritins can also contribute to O₂ detoxification by consuming O_2_ through the ferroxidase reaction (45, 47). DpsA safeguards chromosomal DNA against oxidative and nutrient stress while also functioning as a ferritin-like protein to sequester iron and mitigate reactive oxygen species (45). From a synthetic biology perspective, it is noteworthy that an *E. coli* strain engineered to express the full complement of genes required for N_2_ fixation accumulated its native DpsA protein under N_2_ fixation conditions (48). This observation is particularly significant, as the mechanisms by which a non-diazotrophic host maximizes opportunities to fine-tune the acquired capacity for N_2_-fixation remain largely unexplored.

Elhai (2026) reported that diazoplast genomes, including that of *E. adnata*, encode an alternative respiratory terminal oxidase (a putative type 1 quinol oxidase) alongside cytochrome *c* oxidase. In *Anabaena* sp. PCC 7120, this enzyme is heterocyst-specific and essential for nitrogenase function by shielding nitrogenase from O₂ inhibition (25).

Additionally, a broader protection can be attributed to the notable enrichment of molecular chaperones in the diazoplast proteome. Evidence from *Anabaena* highlights their importance: mutation of either GroEL or Cpn60 proved lethal, while GroEL overexpression not only mitigated oxygen stress but also enhanced tolerance to heat and salt, increased nitrogenase activity under suboptimal temperatures, and reduced protein aggregation. Thus, chaperone-mediated stabilization of nitrogenase extends beyond shielding against O₂ interference to encompass diverse environmental stressors that challenge protein stability during diazotrophic growth (49).

A recent transcriptomic analysis in *R. gibba* showed a similar trend for the most abundant transcripts corresponding to genes involved in N_2_ fixation, stress and molecular chaperones, membrane transport, oxidative PPP, and oxidative phosphorylation (20). High enrichment in molecular chaperones, especially GroEL and DnaK, was shown to be a common feature of endosymbionts with reduced genome size. This could represent a general response to rapid genome evolution and accumulation of deleterious non-synonymous mutations producing aberrant and misfolded proteins towards pseudogenization and gene-loss (50). Nevertheless, such a mechanism and a more specific role in protecting nitrogenase are not mutually exclusive.

Although both diazoplasts (8, 17) and nitroplasts (51) appear to be more active in N_2_ fixation during the light period, light itself does not seem to strongly regulate *nif* gene expression at the transcriptional (20) or proteome (this study) level. This suggests that light influences N_2_ fixation through other mechanisms, likely involving host-symbiont interactions. In particular, daytime catabolism within the diazoplast may support N_2_ fixation by coupling it to the host’s photosynthetic carbon supply (17, 20, 51). Gene expression analysis supports this interpretation and highlights the role of the oxidative PPP pathway in producing reducing equivalents for N_2_ fixation in diazoplasts. A similar pattern was observed in nitroplast (UCYN-A)-containing *B. bigelowii*, supporting a generalized feature of the lifestyle of N_2_-fixing photosynthetic eukaryotes. Here we have shown regulation of acetylene reduction activity by NH₄⁺ supply in the culture medium. Although with a lower level of detail than in our study, partial inhibition of acetylene reduction activity by NH₄⁺ or NO_3_^-^ was also previously determined in *R. gibba* (20, 52).

Changes in gene/protein expression in response to NH_4_^+^ supply in the *E. adnata* culture medium were very prominent in terms of many proteins across most Clusters of Orthologous Groups (COG). This pattern suggests a general and coordinated regulatory commitment primarily toward diazotrophic metabolism. Nevertheless, the variation in protein abundance was relatively modest for most proteins. Modulation of Nif, PPP, and PetH protein abundance to a similar extent in response to NH₄⁺ supplementation further supports the proposed functional link. Importantly, no change was observed in the number of diazoplasts per diatom cell for up to 28 days in cultures periodically supplemented with NH₄⁺. This rules out the possibility that diazoplast loss during the study could bias the interpretation of protein expression analyses. In longer-term experiments (four months), loss of diazoplasts was observed in *Epithemia* spp. cultures maintained in nitrogen-replete medium (8).

In their recent transcriptomic analysis, Abresch and colleagues (2024) pointed out that in *R. gibba*, the most highly expressed gene in all time points and conditions is a hypothetical protein homologous to the *Rippkaea orientalis* sp. PCC 8801 circadian-oscillating polypeptide 23 (COP23) (20). The abundance of the *E. adnata* homologous protein (EAPSB_01376) was not particularly high in our study, representing only 0.1% of the diazoplast proteome as estimated by emPAI analysis. Nevertheless, it was the protein most strongly down-regulated by NH₄⁺ supplementation, which may also be interpreted as activated under NH₄⁺ deficiency. COP23 was primarily studied as a circadian oscillating protein located in the cell membrane of *Synechococcus/Cyanothece* RF-1 (36), which fixes N_2_ primarily in the dark. In this free-living bacterium, COP23 is regulated by a circadian rhythm protein synthesis/degradation rhythm, the latter being dependent on light and Ca²⁺ (53). COP23 transcript levels also changed more than 5-fold in two different UCYN-A strains (nitroplasts) between the light and dark periods (54). Another proteomic analysis conducted in the N_2_-fixing cyanobacterium *Anabaena* sp. PCC 7120 uncovered a highly expressed protein (Alr2313), homologous to COP23, whose abundance increased 3.6-fold or 6.7-fold after 3 or 12 h after removing combined nitrogen from the culture medium (55). Although available evidence suggests a possible involvement of COP23 in the regulation of N_2_ fixation, its specific function and putative regulatory role during diazotrophy, whether in free-living diazotrophic bacteria or in diazoplasts/nitroplasts, remains unresolved. Moreover, the possibility that COP23 is regulated by intracellular 2-oxoglutarate levels, sensing the cellular C/N ratio through NH₄⁺ assimilation via the GS–GOGAT cycle and signaling through NtcA to activate Class II promoters, opens an appealing avenue for future studies of diazoplast regulation. The GS–GOGAT–PII/GlnK–NtcA–PipX axis, possibly involving Amt1 for ammonium transport and sensing, would allow diazoplasts to perceive the nitrogen status of the host, whether in the form of NH₄⁺ itself or NH₄⁺ derived from host nitrate assimilation. In parallel, host-supplied carbon modifies the C/N ratio within the diazoplast, triggering proteomic modulation toward achieving nitrogen homeostasis at the holobiont level.

In free-living N₂-fixing cyanobacteria, cyanophycin granules function as dynamic nitrogen reserves to balance assimilated carbon and nitrogen, thereby sustaining growth under fluctuating conditions (34). The absence of cyanophycin metabolism in diazoplasts is noteworthy as it removes this autonomous safeguard, rendering nitrogen balance dependent on host-derived carbon and nitrogen fluxes. This loss underscores a shift from cell-level regulation to holobiont-level integration.

For synthetic biology, this has profound implications: in a putative transgenic plant engineered to express *nif* genes within organelles, successful N_2_ fixation would depend not only on introducing the enzymatic machinery but also on establishing regulatory circuits that integrate organelle activity with host C/N fluxes. Dynamic sensing of NH_4_^+^ and C availability, coupled with feedback control of reductant allocation and metabolite transport, would be essential to achieve optimal nitrogen supply and stable C/N homeostasis in engineered crops.

In summary, our findings considerably advance understanding of the makeup and regulation of N₂ fixation in eukaryotes. By integrating genomic and proteomic evidence, we reinforce the view that diazoplasts are metabolically committed to N₂ fixation, requiring coordinated ATP and NADPH production along with accessory functions that sustain this process in the context of an endosymbiotic lifestyle. Our results highlight the likely-roles of PPP and PetH in supplying reductant to NifH, of plasmid-encoded GlpF for glycerol transport as a key mechanism of metabolic integration, and a concerted strategy to cope with oxidative stress, which could have been crucial factors at the onset of the evolutionary pathway from ancestral to extant symbiotic N₂-fixing diatoms.

Beyond their evolutionary significance and contribution to understanding organogenesis pathways, these results hold translational implications. By unveiling the metabolic specialization, integration, and regulation strategies of diazoplasts and their hosts, our study provides a framework for ongoing efforts to engineer N₂-fixing capabilities into agricultural plants (56, 57). Such advances could reshape biogeochemical cycles and contribute to sustainable agriculture by reducing dependence on synthetic nitrogen fertilizers.

## Materials and Methods

### Cell isolation, visualization and culture conditions

*Epithemia adnata* LB21 was recovered from the periphyton of the macrophyte *Schoenoplectus californicus* collected from the littoral zone in La Brava shallow lake, Buenos Aires, Argentina (37°52′52.1″ S, 57°58′38.3″ W) in July 2021 and March 2022. Individual cells were isolated using streaking-plate and micropipetting techniques. Established strains are cultured in FB-N medium (*SI Appendix*, additional materials and methods) lacking a N source other than N_2_ from the air and incubated at 21°C under a 12:12 h light:dark cycle at 50 μm photons · m^-2^ · s^-1^ provided by 16-Watt LED tubes (LEDVANCE cat. #7019143). Cultures were routinely maintained in 1.5% (w/v) agar slants and liquid cultures were used for most experiments. Alternatively, cultures were long-term preserved in FB-N/agar at 4 °C in a translucent refrigerator to allow ambient light (indoors) exposure for 15 months. For growth analyses, 5-30 diatom cells were transferred by micropipetting into 24 multiwell racks containing FB-N/agar medium. When indicated, NH_4_Cl or Ca(NO_3_)_2_ was supplemented at the stated concentrations. Cells were counted daily using a stereomicroscope (Nikon SMZ800).

For brightfield and epifluorescence microscopy imaging, cells were fixed in 4% (v/v) paraformaldehyde in FB-N medium for 30 min and centrifuged at 100 × *g* for 2 min. Pelleted cells were washed twice, resuspended FB-N medium containing 10% (v/v) DMSO and 1× SYBR Green I (FMC BioProducts, Rockland, ME, USA, Cat. #50513) and incubated in the dark for 15 min. Rinsed cells were mounted in 90% (v/v) glycerol, 0.1× PBS, and 1 mg/mL p-phenylenediamine, pH 8.0. Micrographs were captured using an Olympus DP72 digital camera mounted on a Nikon Eclipse E600 microscope at 1000 × magnification. Epifluorescence imaging was performed using Nikon B2-A and G-2E/C filter cubes for SYBR Green I and chlorophyll, respectively, and images were acquired using Olympus cellSens Entry software (v.1.6). For scanning electron microscopy, cell organic matter was oxydized and removed by boiling for one hour in 30% (v/v) hydrogen peroxide, rinsed four times with distilled water, and left to stand overnight in 10% (v/v) HCl. This procedure was repeated twice. Pelleted frustules were resuspended in distilled water, and 10 µL samples were allowed to dry in an oven at 50 °C overnight onto 10 × 10 mm glass holders, and coated with chromium. Micrographs were acquired using a Zeiss Crossbeam 350 Field Emission Scanning Electron Microscope (FE-SEM) equipped with EDS, EBSD, and FIB at the Laboratory of Electron Microscopy of the Institute of Materials Science and Technology Research (LAMEI-INTEMA).

### Acetylene Reduction Assay (ARA)

For ARA, 4 mL samples of *E. adnata* culture were dispensed into 12-mL glass vials capped with aluminum foil. At 12pm and 12am, for two consecutive days, six vials were crimp-sealed with a butyl rubber stopper. A final concentration of 10% (v/v) acetylene was obtained by replacing 750 µl of headspace with an equivalent volume of pure acetylene. Vials were disposed thumbed and incubated with agitation at 150 rpm on an orbital shaker under the same growth conditions, including the light or darkness phases of the diel cycle for one hour. After withdrawing 1 mL of culture for protein determination, reactions were stopped by adding 240 µL of 17.8% (w/v) paraformaldehyde (CAS No 30525-89-4) and 760 µL of sterile FB-N media (for pressure equilibration). A gas sample of 100 µL from the vial headspace was injected into a Shimadzu Nexis GC-2030 gas chromatograph coupled with a flame ionization detector and an SH-AluminaBOND/Na_2_SO_4_ PLOT column (30 m x 0.55 mm, 10 µm of film thickness). The injector was maintained at 200 °C, and the column was programmed with a temperature gradient from 90 °C to 120 °C over 3 min; N_2_ was used as a carrier gas at 15 mL · min^-1^. Data collection, peak identification and ethylene quantification were performed using LabSolutions (v5.111) software, and compared to a standard curve prepared with pure ethylene. Whole protein of the samples was determined by the Lowry method, after addition of NaOH at 1N final concentration.

When indicated, diatoms were acclimated to growth in the presence of NH_4_^+^ (250 µM NH_4_Cl and 500 µM HEPES/KOH, pH 7.5) for 15 days with complete medium replacement every ∼72 h throughout the experiments. Then cells were switched to the indicated conditions for ARA determination. Diazotrophic cultures were also supplemented with HEPES/KOH. For the determination of the NH_4_^+^ remaining in the culture medium, 3.0 mL of supernatant, after stopping the ARA determinations was filtered with 0.22 µM polyvinylidene fluoride membrane and NH_4_^+^ was determined by the indophenol method essentially as reported before (58).

### Immunoblot analysis

Frozen pellets stored at −80 °C (∼100 µL) were resuspended in 2× ice-chilled buffer (final concentration: 25 mM Tris-HCl, pH 7.5; 50 mM NaCl; 0.5 mM PMSF) at a 1:1.5 (sample:buffer) volume ratio. Cells were disrupted by two cycles of sonication (35% amplitude, 15 pulses of 7 s ON/10 s OFF) on ice, followed by centrifugation at 13,000 x *g* for 15 min at 4 °C. The supernatant was collected, and protein concentration was determined using the bicinchoninic acid assay (BCA; Thermo Scientific, Cat. No. 23223) with bovine serum albumin as the standard. Protein samples (10 - 30 µg) were subjected to SDS-PAGE, transferred to nylon membranes and incubated with antibodies against the *Azotobacter vinelandii* NifH, NifDK, NifEN, or NifB and incubated overnight at 4 °C with gentle orbital shaking. Immuno signals were developed using goat anti-rabbit IgG antibodies conjugated to alkalyne phosphatase (Sigma Aldrich, Cat. No. A8025) and finally incubated in the presence of 5-bromo-4-chloro-3-indolyl phosphate and nitro blue tetrazolium chloride, essentially as described before (58). Densitometry signals were acquired using the Fiji (ImageJ) software and expressed as arbitrary units.

### Genomics

For DNA extraction, diatom samples were recovered from the surface of FB-N/agar slants (R1 sample) and from FB-N liquid cultures (R2 sample). Because diatom cultures produce copious amount of exopolysaccharides, for its partial removal, biomass was resuspended in 500-1000 µL of sterile FB-N medium and gently sonicated (35 s, 30% amplitude; 5 s ON/10 s OFF; 2 cycles) in an ice-water bath, followed by centrifugation at 100 × *g* for 2 min at 4°C. Approximately 100 µL of biomass was recovered and washed with sterile TE buffer and centrifuged at 12,000 × g for 5 min at 4°C. DNA was extracted using chloroform–phenol as described (59).

For long-read Oxford Nanopore sequencing (ONT), DNA libraries were prepared using Ligation Sequencing Kit V14 (SQK-LSK114) according to the manufacturer’s instructions and sequenced on a MinION flow cell R10.4.1 at the Centre for Plant Biotechnology and Genomics (CBGP, UPM-INIA/CSIC), Madrid, Spain. Basecalling was performed using Guppy (v3.5.2) (ONT). FASTQ file quality was checked with FastQC (v.0.11.9), and adapter trimming and read filtering were performed using PoreChop (v.0.2.4) (https://github.com/rrwick/Porechop) and Chopper (v.0.6.0) and a total of 1,397,366 long-reads were obtained (Accession: SRR37659450). For Illumina sequencing, two independent short-read libraries were prepared using the Nextera DNA Flex Library Prep Kit (Illumina). Multiplexed paired-end sequencing (2 × 250 bp) was performed on a NovaSeq 6000 platform at the Genomics Core Facility of the Andalusian Centre for Molecular Biology and Regenerative Medicine (CABIMER). Raw reads from samples R1 and R2 (Accessions: SRR37659451, SRR37659452, SRR37659453, and SRR37659454) from different flow cells were concatenated, resulting in 24,558,678 and 22,016,821 reads, respectively. Redundant sequences were then collapsed into single representatives using CD-HIT-EST (v. 4.8.1). Read quality was assessed with FastQC (v. 0.11.9). De novo assembly of paired-end reads was performed using MEGAHIT assembler (v.1.2.9), and contigs shorter than 1 kb were discarded. Quality profiles and basic statistics of assemblies were generated with QUAST (v.5.2.0).

For the assembly of the *E. adnata* diazoplast genome, a hybrid approach was followed using both short and long sequencing reads and the SAMBA scaffolder from MASURCA toolkit (v.4.1.0). A 2,782,194 nt circular genome was annotated with Prokka (v1.14.6) and eggNOGmapper (v2.1.12). Assembly completeness based on essential genes was assessed using BUSCO (v5.5.0) with cyanobacteria_odb10 database.

Prodigal (v.2.6.3) was employed to perform protein prediction, and amino acid sequences were aligned to a homemade protein reference database by using Diamond (v2.1.6.160) BLASTP in ultra-sensitive mode, with the default scoring matrix. The reference database was built from: eggNOG full database (v5.0), EukProt full database (v03.2021_11_22), custom NCBI genomes and other sequences coming from diatom endosymbionts or phylogenetically close free-living cyanobacteria (mostly *Crocosphaera* sp. and *Cyanothece* sp.) and Custom NCBI genomes and other sequences from diatoms. Average nucleotide identity (ANI) was estimated and a UPGMA tree was constructed among genomes of known diazoplasts using OrthoANI.

Nif-proteins synteny diagram was constructed using Simple Synteny, with the portion of genomic sequence involving gene cluster from *nifT* and *fdxB* genes of all known spheroid bodies. Diagrams were customized using Inkscape. Details and references for methods for diazoplast and diatom mitochondria and chloroplast genomes assembly, annotation and analysis are presented in *SI Appendix*, EapSB genome assembly.

### Proteomics

Protein extraction and quantification were carried out as described above for immunoblot analysis. Sample proteins in 1x Laemmli buffer were briefly resolved (1 cm) in a 0.75-mm-thick 12% polyacrylamide gel. Gels were fixed for 45 min in a fixing solution (30% (v/v) methanol, 20% (v/v) acetic acid) and stained for 5 min in Colloidal Coomassie Blue (0.12% (w/v) Coomassie Brillant Blue G-250, 10% (v/v) H₃PO₄, 10% (w/v) (NH_4_)_2_SO_4_; 20% (v/v) methanol). Protein bands were excised and submitted to the Mass Spectrometry Unit of the Institute of Molecular and Cellular Biology of Rosario (UEM-IBR), Argentina for LC-MS/MS proteomic analysis. Proteome Discoverer (v2.4.1.15) was used for protein identification and quantification and Perseus (v1.6.15.0) for statistical analysis. Protein relative abundances were estimated by exponentially modified protein abundance index (emPAI) analysis (34). Additional details on proteomics analyses are presented as *SI Appendix*, Proteomics: Mass spectrometry data analysis. The mass spectrometry proteomics data have been deposited to the ProteomeXchange Consortium via the PRIDE (60) partner repository with the dataset identifier PXD077509.

### Data, Materials, and Software Availability

All study data are included in the article and/or *SI Appendix*.

## ACKNOWLEDGMENTS

We acknowledge valuable technical assistance from Drs. Marisol Fassolari and Macarena Perez Cenci for assistance for GC analysis. This study was supported by the National University of Mar del Plata Grant 80020240500156MP to L. C. and Gates Foundation Grant INV-067006 to L. M. R.

## Author contributions

1. L. C. designed the research; A. O. S., I. A., M. D. N., L. S. R., A. S. M., A. L. G., S. G.B., and L. C. performed research; E. F. handled sequencing samples; all authors analyzed the data and edited the draft manuscript; L. M. R. and L. C. acquired funding for the research; L. C. wrote the draft manuscript; A. O. S. prepared the manuscript figures.

