## Supplementary Material/Information for "Metabolic commitment and nitrogen control of diazotrophy in the diazoplast-containing diatom *Epithemia adnata*"

*SI Appendix*

### Supplementary materials and methods

#### Diatom culture medium

The FB-N culture medium was originally developed by Floener and Bothe, later modified by Adler et al. (2010), and further adapted in our study. It contained: 0.23 mM $CaSO_4 \cdot 2\ H_2O$; 1.20 mM KCl; 2.5 mM NaCl; 1.00 mM $MgSO_4 \cdot 7\ H_2O$; 0.38 mM $K_2HPO_4 \cdot 3\ H_2O$; 1.8 mM $Na_2SiO_3 \cdot 9\ H_2O$; 4.00 mM $NaHCO_3$ (macronutrients). 100.00 µM $Na_2EDTA \cdot 2\ H_2O$; 0.50 µM $MnCl_2 \cdot 4\ H_2O$; 0.12 µM $FeCl_3 \cdot 6\ H_2O$; 5.10 µM $NaMoO_4 \cdot 2\ H_2O$; 0.05 µM $CuSO_4 \cdot 5\ H_2O$; 0.09 µM $CoCl_2$; 0.05 µM $ZnSO_4 \cdot 7\ H_2O$; 0.05 µM $NaB_4O_7$ (micronutrients).

L. Floener, H. Bothe, Nitrogen fixation in *Rhopalodia gibba*, a diatom containing bluegreenish inclusions symbiotically. In: Schwemmler W, Schenk H. (eds) Endocytobiology: Endosymbiosis and Cell Biology, a Synthesis of Recent Researc , Vol. 1, Walter de Gruyter & Co. Berlin, pp. 541-552 (1980).

S. Adler, L. M. Derch, Maier, U. G., Cultivation of the diatom *Rhopalodia gibba*. *Endocytobiosis Cell. Res.* **20** (2010).

#### EapSB genome assembly

A hybrid assembly was generated from Illumina primary contigs and ONT reads using the SAMBA scaffolder within the MaSuRCA toolkit (v4.1.0). The two final scaffolds obtained suggested overlap, resulting in two possible draft genome models. A gap of ~6 kb was inferred by comparison with the EtSB genome (AP012549.1), which in all known SB genomes contains a second ribosomal gene cluster (16S, Ile-tRNA, 23S, and 5S in tandem). The two draft models were manually joined and annotated using Prokka (v1.14.6). Terminal genes were considered as flanking markers of the ribosomal cluster, and ONT reads containing ribosomal gene copies were identified using megablast (e-value = $1e^{-7}$). ONT reads >10 kb were retained and annotated with Prokka. Manual inspection of annotated reads revealed three ONT sequences consistent with one of the draft models based on synteny, while no ONT support was found for the alternative model. Identified ONT reads were aligned with MUSCLE (default parameters) and visualized in Jalview (v2.11.4.0), yielding a consensus sequence of 24,764 bp. Two paired-end Illumina datasets were mapped to this consensus using Bowtie2 (v2.5.3), and error correction was performed with LoRDEC (v0.9, default parameters). The corrected consensus sequence was manually trimmed to close the EapSB genome gap. The resulting circular chromosome was annotated using Prokka (v1.14.6) and the eggNOGMapper web tool. All analyses were performed on the Galaxy Europe platform (Galaxy Europe platform).

**Diazoplastic plasmid, mitochondrial and chloroplast genomes**

ONT long-reads were mapped using Bowtie2 (v.2.5.3) to diazoplastic plasmid sequence (CP076463.1), chloroplast sequences (OR527427.1, OR527428.1, OR527429.1), and mitochondrial sequence (OR527426.1) of E. adnata 19Bon2 used as a reference. Putative diazoplastic plasmid and chloroplast ONT-reads were assembled using Flye (v2.9.5) with three cycles of polishing. Five rounds of polishing were further performed using Racon (v1.5.0) and Medaka (v2.1.1) (basecaller model: r1041_e82_400bps_hac_g632) with ONT long-reads, followed by Polypolish (v0.6.1) and Pilon (v1.2) with Illumina short-reads. Putative mitochondrial ONT were mapped with one set of paired-end Illumina reads for error correction using Lordec (v0.9). Another set of Illumina was mapped to mitochondrial reference sequence. Corrected ONT and the second set of mapped short reads were hybrid assembled using Unicycler (v0.5.1). Mapping steps were performed using Minimap2(v2.28).

The complete analysis was conducted on the Galaxy Europe platform. Complete circular chromosomes were obtained of 5,640 bp for diazoplast's plasmid, and 120,460 bp for chloroplast genome, and 36,719 bp for mitochondrial genome. Diazoplastic plasmid and mitochondrial sequences were annotated using Prokka (v1.14.6), whereas the plastid genome were annotated using MFannot (https://megasun.bch.umontreal.ca/apps/mfannot/).

**References to software and databases**


CD-HIT-EST (v. 4.8.1) - W. Li, A. Godzik, Cd-hit: a fast program for clustering and comparing large sets of protein or nucleotide sequences. *Bioinformatics* (Oxford, England) **22**, 1658–1659 (2006). https://doi.org/10.1093/bioinformatics/btl158.

MEGAHIT assembler (v.1.2.9) - D. Li, C. M. Liu, R. Luo, K. Sadakane, T. W. Lam, MEGAHIT: an ultra-fast single-node solution for large and complex metagenomics assembly via succinct de Bruijn graph. *Bioinformatics* (Oxford, England) **31**, 1674–1676 (2015). https://doi.org/10.1093/bioinformatics/btv033.

QUAST (v.5.2.0) - A. Gurevich, V. Saveliev, N.Vyahhi, G. Tesler. QUAST: quality assessment tool for genome assemblies. *Bioinformatics* (Oxford, England) **29**, 1072–1075 (2013). https://doi.org/10.1093/bioinformatics/btt086.

MASURCA toolkit (v.4.1.0) - A. V. Zimin, G. Marçais, D. Puiu, M. Roberts, S. L. Salzberg, J. A. Yorke, The MaSuRCA genome assembler. *Bioinformatics* (Oxford, England) **29**, 2669–2677 (2013). https://doi.org/10.1093/bioinformatics/btt476.

Prokka (v1.14.6) - T. Seemann, Prokka: rapid prokaryotic genome annotation. *Bioinformatics* (Oxford, England) 30, 2068–2069 2014. https://doi.org/10.1093/bioinformatics/btu153.

eggNOGmapper (v2.1.12) - J. Huerta-Cepas, et al., eggNOG 4.5: a hierarchical orthology framework with improved functional annotations for eukaryotic, prokaryotic and viral sequences. *Nucleic Acids Res.* **44**(D1), D286-D293 (2016).

BUSCO (v5.5.0) - M. Seppey, M Manni, E. M. Zdobnov, BUSCO: Assessing Genome Assembly and Annotation Completeness. *Methods Mol. Biol.* (Clifton, N.J.), **1962**, 227–245 (2019). https://doi.org/10.1007/978-1-4939-9173-0_14.

Prodigal (v.2.6.3) - D. Hyatt, et al., Prodigal: prokaryotic gene recognition and translation initiation site identification. *BMC bioinformatics* **11**, 119 (2010). https://doi.org/10.1186/1471-2105-11-119.

Diamond (v2.1.6.160) - B. Buchfink, H. Ashkenazy, K. Reuter, J. A. Kennedy, H. G. Drost, "Sensitive clustering of protein sequences at tree-of-life scale using DIAMOND DeepClust". *bioRxiv* 2023.01.24.525373 (2023). https://doi.org/10.1101/2023.01.24.525373

eggNOG full database (v5.0) - J. Huerta-Cepas, et al., eggNOG 5.0: a hierarchical, functionally and phylogenetically annotated orthology resource based on 5090 organisms and 2502 viruses. *Nucleic Acids Res.* **47**(D1), D309–D314 (2019). https://doi.org/10.1093/nar/gky1085.

EukProt full database (v03.2021_11_22) - D. J. Richter, et al., EukProt: A database of genome-scale predicted proteins across the diversity of eukaryotes. *Peer Community J.* **2**, e56 (2022).

OrthoANI - I. Lee, Y. Ouk Kim, S. C. Park, J. Chun, OrthoANI: An improved algorithm and software for calculating average nucleotide identity. *Int. J. Syst. Evol. Microbiol.* **66**, 1100–1103 (2016). https://doi.org/10.1099/ijsem.0.000760.

MUSCLE - R. C. Edgar, MUSCLE: multiple sequence alignment with high accuracy and high throughput. *Nucleic Acids Res*. **32**, 1792-1797. (2004).

Jalview (v2.11.4.0) - A. M. Waterhouse, J. B. Procter, D. M. A. Martin, M. Clamp, G. J. Barton, Jalview Version 2—a multiple sequence alignment editor and analysis workbench. *Bioinformatics* **25**, 1189–1191 (2009). https://doi.org/10.1093/bioinformatics/btp033.

Bowtie2 (v2.5.3) **–** B. Langmead, S. L. Salzberg, Fast gapped-read alignment with Bowtie 2. *Nat. Methods* **9**, 357–359 (2012). https://doi.org/10.1038/nmeth.1923.

LoRDEC (v0.9) – L. Salmela, E. Rivals, LoRDEC: Accurate and efficient long read error correction. *Bioinformatics* 30, 3506–3514 (2014). https://doi.org/10.1093/bioinformatics/btu538.

Minimap2 (v2.28) – H. Li, Minimap2: Pairwise alignment for nucleotide sequences. *Bioinformatics*, **34**, 3094–3100 (2018). https://doi.org/10.1093/bioinformatics/bty191.

Flye (v2.9.5) – Y. Lin, J. Yuan, M. Kolmogorov, M. W. Shen, M. Chaisson, P. A. Pevzner, Assembly of long error-prone reads using de Bruijn graphs. *Proc. Natl. Acad. Sci. U.S.A.* **113**, E8396–E8405 (2016). https://doi.org/10.1073/pnas.1604560113.

Unicycler (v0.5.1) – R. R. Wick, L. M. Judd, C. L. Gorrie, K. E. Holt, Unicycler: Resolving bacterial genome assemblies from short and long sequencing reads. *PLoS Comput. Biol.* **13**, e1005595 (2017). https://doi.org/10.1371/journal.pcbi.1005595.

Racon (v1.5.0) – R. Vaser, I. Sović, N. Nagarajan, M. Šikić, Fast and accurate de novo genome assembly from long uncorrected reads. *Genome Res*. **27**, 737–746 (2017). https://doi.org/10.1101/gr.214270.116.

Medaka (v2.1.1) – Ltd., O. N. T. (2020). medaka. https://github.com/nanoporetech/medaka.

Polypolish (v0.6.1) – R. R. Wick, K. E. Holt, Polypolish: Short-read polishing of long-read bacterial genome assemblies. *PLoS Comput. Biol.* **18**, e1009802 (2022). https://doi.org/10.1371/journal.pcbi.1009802.

Pilon (v1.2) – B. J. Walker, T. Abeel, T. Shea, M. Priest, A. Abouelliel, S. Sakthikumar, C. A. Cuomo, Q. Zeng, J. Wortman, S. K. Young, A. M. Earl, Pilon: An Integrated tool for comprehensive microbial variant detection and genome assembly improvement. *PLoS One* **9**, e112963 (2014). https://doi.org/10.1371/journal.pone.0112963.

**LC-MS/MS analysis**

An in-gel trypsin digestion was carried out according to Link and LaBaer (2009). Three microliters of protein sample were loaded on a nanoHPLC Ultimate3000 (Thermo Scientific) for peptide separation using a 50 cm C18 nano column (µPAC Neo, Thermo). The mobile phase flow rate was 300 nL · min using 0.1% formic acid in water (solvent A) and 0.1% formic acid in acetonitrile (solvent B). The gradient profile was set as follows: 4-30% solvent B for 64 min, 30-80% solvent B for 7 min and 80% solvent B for 1 min. The mass spectra were obtained using a Thermo Scientific Q-exactive HF. For ionization, 1.9 kV of liquid junction voltage and 300°C of capillary temperature were used. The full scan method employed a m/z 375–2,000 mass selection, an Orbitrap resolution of 120,000 (at m/z 200), a target automatic gain control (AGC) value of $1 \cdot e^6$ and a maximum injection time of 100 ms. After the survey scan, the top 15 of most intense precursor ions were selected for MS/MS fragmentation. Fragmentation was performed with a normalized collision energy of 28 eV and MS/MS scans were acquired with a dynamic first mass. The AGC target was $5\ e^5$, resolution of 30,000 (at m/z 200), intensity threshold of $1.5 \cdot e^5$, isolation window of 1.4 m/z units and maximum injection time of 55 ms. Charge state screening was enabled to reject unassigned, singly charged, and equal or more than six protonated ions. A dynamic exclusion time of 30 sec was used to discriminate against previously selected ions.

J. Link, J. LaBaer, In-gel trypsin digest of gel-fractionated proteins. Cold Spring Harbor Protocols, 2009, pdb-prot5110 (2009).

**Proteomics: Mass spectrometry data analysis**

MS data were analyzed using Proteome Discoverer (v2.4.1.15) with standardized workflows. Raw mass spectra (*.raw) files were searched against a database of predicted proteins derived from partial nuclear and complete extranuclear genomes. Precursor and fragment mass tolerances were set to 10 ppm and 0.02 Da, respectively, allowing up to two missed cleavages. Dynamic modifications were restricted to a maximum of four per peptide, with no more than three identical modifications per peptide. Variable modifications included methionine oxidation, N-terminal acetylation, and methionine loss (with or without acetylation). Carbamidomethylation of cysteine was specified as a static modification.
The Proteingroups.txt output from Proteome Discoverer was analyzed in Perseus (v1.5.16.0). Protein abundance values were $log_2$-transformed for normalization Only proteins identified with more than two or more unique peptides and present in at least 75% of samples of at least one per condition were retained for downstream analysis. Missing values were imputed from a normal distribution of the total matrix (width = 0.3; down-shift = 1.8). Comparisons among conditions were assessed using a two-sided Student’s t-test with a false discovery rate (FDR) of 0.05 and 250 randomizations. The resulting matrix was annotated using eggNOG Mapper and Prokka output files for diazoplast proteins.

For emPAI analysis, raw files were converted to MGF format and processed with Mascot (v2.7.0) using the same fixed and dynamic modification settings, a peptide mass tolerance of 10 ppm, a fragment mass tolerance of 20 ppm, and a maximum of two missed cleavages. Searches were performed against the predicted protein database of the *E. adnata* diazoplast. Output tables were exported, and plots were generated using GraphPad Prism (v9.0.0).

**Supplementary results**

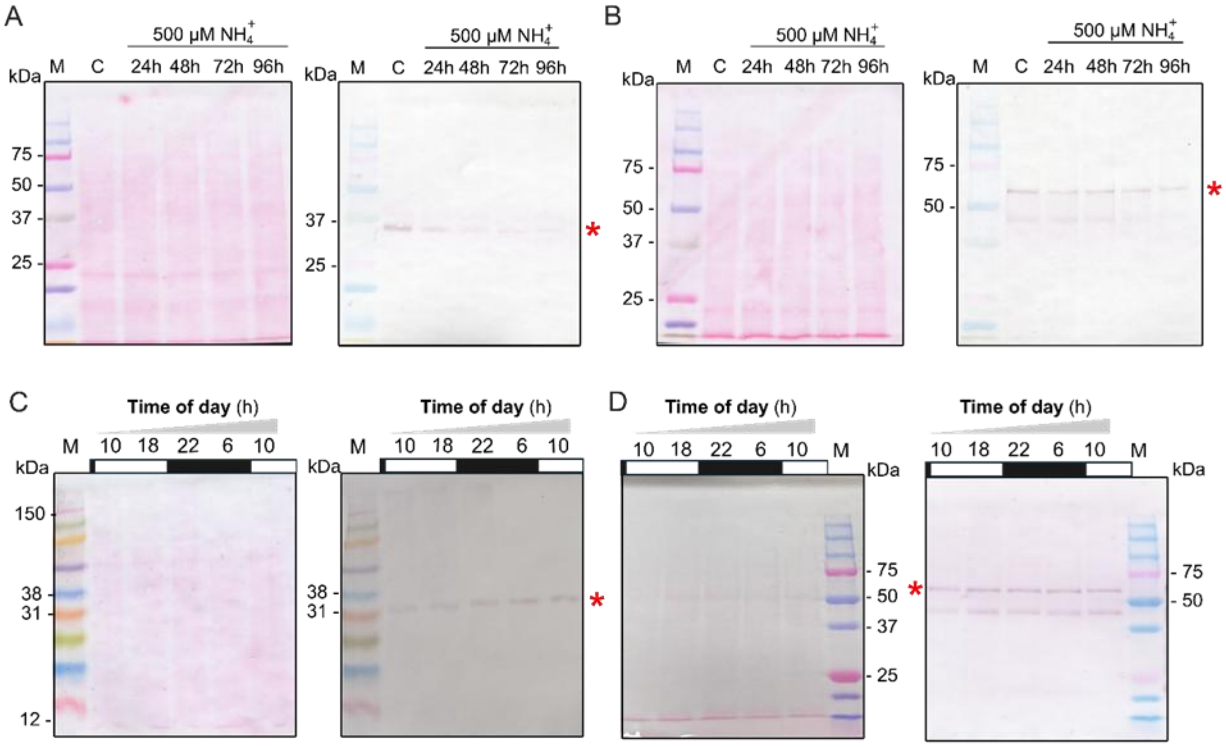


**Fig. S1.** Immunoblot analysis of NifH and NifDK expression in *E. adnata* diazoplasts. Time course expression of *(A)* NifH and *(B)* NifK in *E. adnata* cultures for four consecutive days after $NH_4^+$ supplementation, with samples collected at the midday of the subjective photoperiod. Time course expression of *(C)* NifH and *(D)* NifK in *E. adnata* diazotrophic cultures at different time points of a subjective day (6 and 10 hours correspond to 6 am and 10 am; and 18 and 22 h to 6 pm and 10 pm of the subjective photoperiod). White or black bars indicate the light or dark, respectively. * Indicate the position of the NifH or NifK reactive signals. The subpanel at the left side of each panel represents a protein loading control after staining the nitrocellulose filters with the Ponceau S dye.

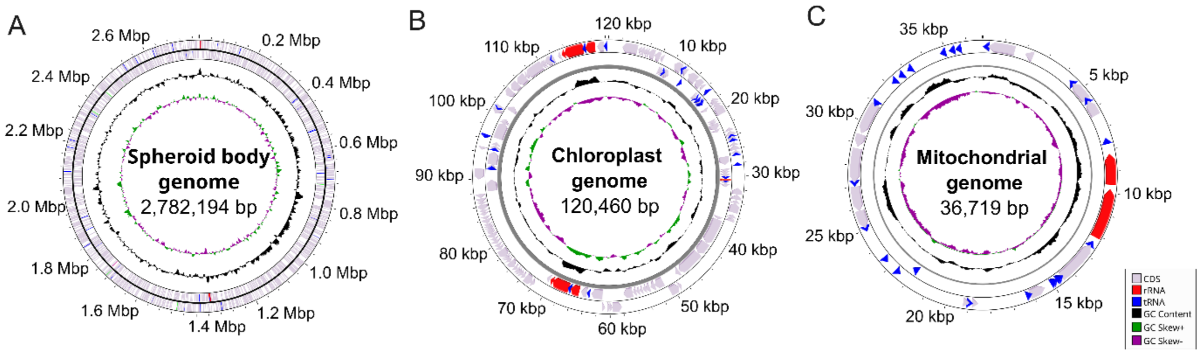


**Fig. S2.** Physical maps of *E. adnata* (*A*) diazoplast (spheroid body), (*B*) chloroplast, and (*C*) mitochondrial genomes. Maps were generated using Proksee (https://proksee.ca/) and edited in Inkscape(v1.4).

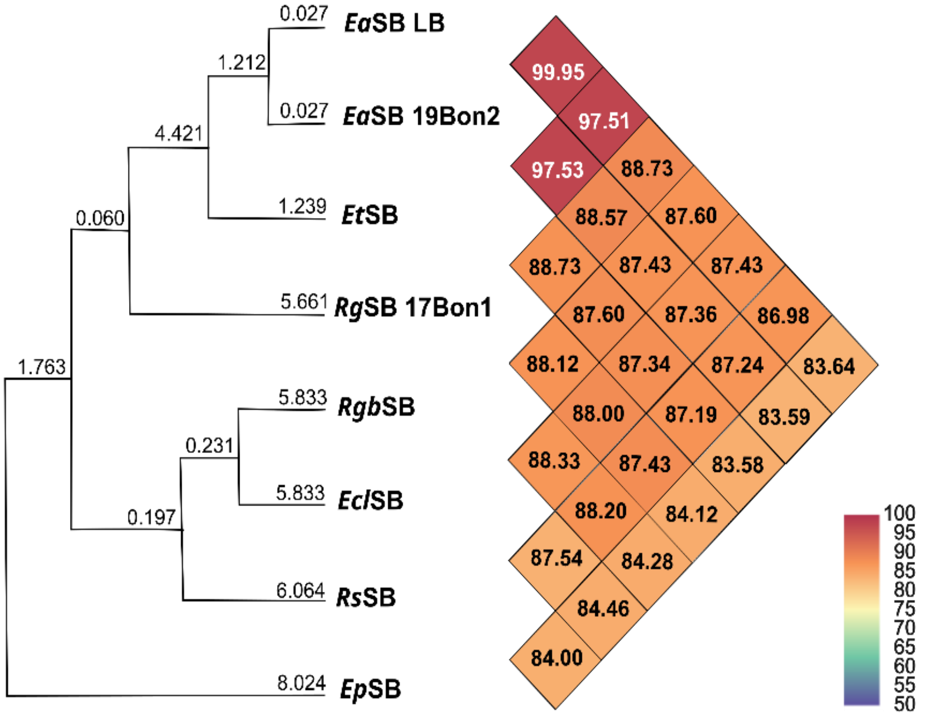


**Fig. S3.** OrthoANI analysis of known diazoplast genomes. Diazoplast genome sequence accession: *E. adnata* isolate LB21 (EaSB LB, this study: ), *E. adnata* isolate 19Bon2 (EaSB 19Bon2: CP076462.1), *E. turgida* (EtSB: AP012549.1), *R. gibba* isolate 17Bon1 (RgSB 17Bon1: CP067995.1), *R. gibberula* (RgbSB: NZ_AP018341.1 ), *E. clementina* (EclSB: CP123991.1), *R. sterrenburgii* (RsSB: CP192514.1), *E. pelagica* (EpSB: NZ_OX373462.1).

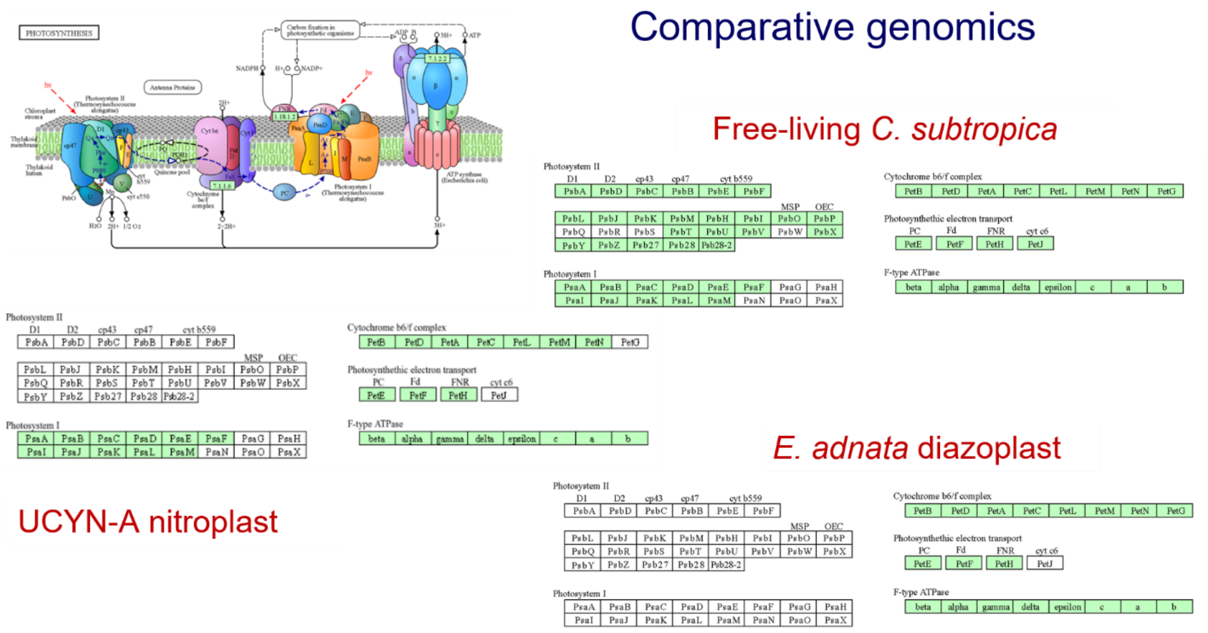


**Fig. S4.** KEGG analysis of predicted photosynthetic genes in the *E. adnata* diazoplast genome.

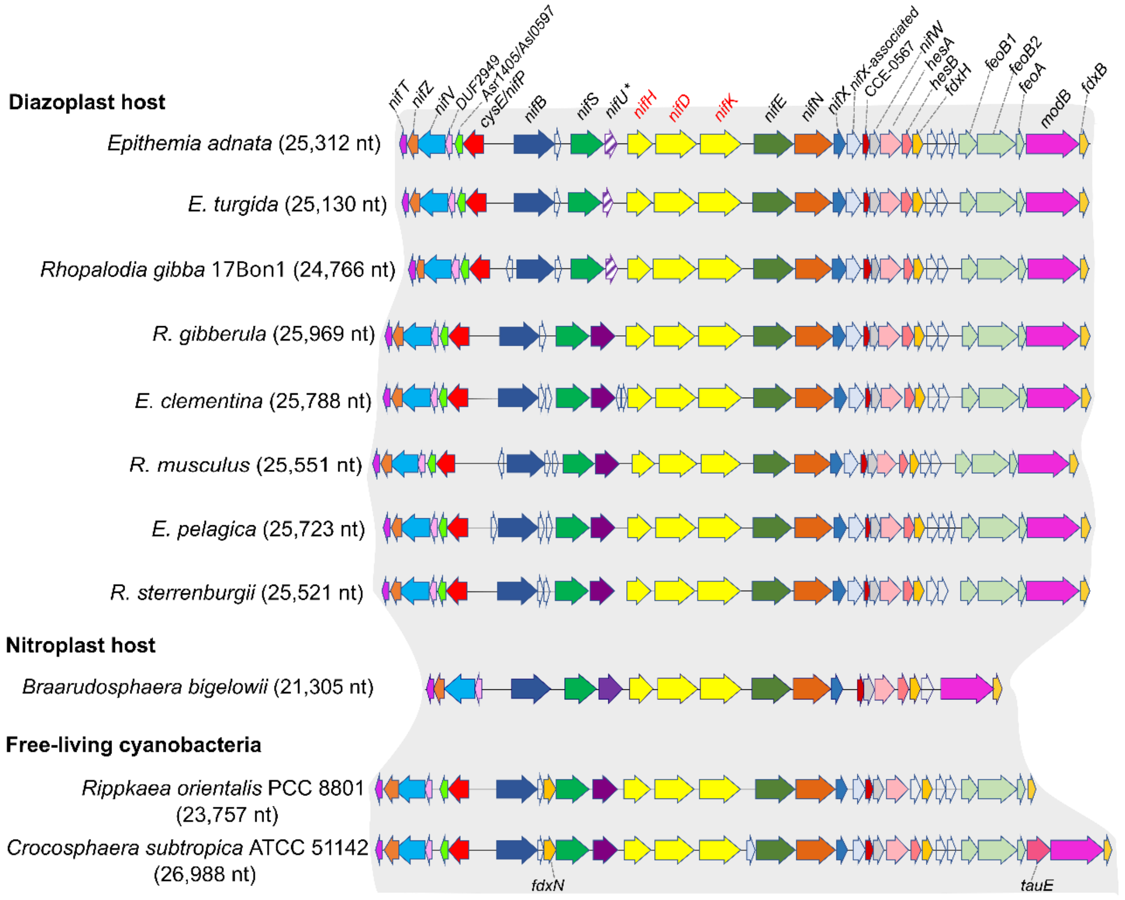


**Fig. S5.** Syntheny of the *nif*-genes clusters in known diazoplasts, and nitroplast, and closed related cyanobacteria. Open boxes or arrows represent hypothetical proteins. The full-length NifU is shown as a solid purple box or arrow, whereas a shortened, purple-hatched version indicates a pseudogenized form.

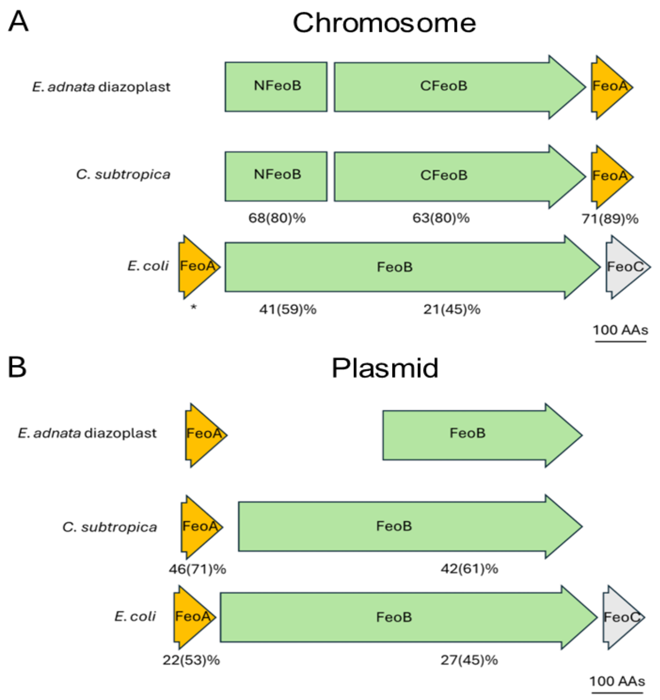


**Fig. S6.** Schematic of *feo* gene arrangements in the *E. adnata* diazoplast genome. Comparison of *feoAB* gene cluster assembly and sequence with those *C. subtropica* ATCC 51142, as a predicted relative to a free-living ancestor of diazoplasts and those of *E. coli*, as a thoroughly studied reference model sequence-structure and functional studies. *(A)* FeoBA copy in the chromosome of *E. adnata* diazoplast physically associated to the *nif* genes locus. *(B) E. adnata* FeoAB copy encoded in the diazoplast plasmid. Numbers under the gene models indicated % identity and homology between parenthesis after BLAST analysis using the *E. adnata* FeoAB sequences as query. * in *(A)* indicates no significant homology found using *E. adnata* FeoA. A similar analysis indicated a similarity of 28(47)% between *C. subtropica* and *E. coli* FeoA. *Epitemia adnata* FeoB copies are 28 (47)% identical, while FeoA copies are even more distantly related. As demonstrated by Gómez-Garzón et al., (2022), the phylogenetic relationship among cyanobacterial FeoA-like proteins is puzzling.

C. Gómez-Garzón, J. E. Barrick, S. M. Payne, Disentangling the evolutionary history of Feo, the major ferrous iron transport system in bacteria. *MBio* **13**, e03512-21. (2022).

```
E.adnata       MSKQTKLSSEVLAEGVGTFVLIFIGTGTIMVNTITEGAITHFGICVVFGAIVTAIIYTIG  60
               MS+ + L  + +AE +GT +LIF G G +    +    +  + I V++G  V   IY
E.coli         MSQTSTLKGQCIAEFLGTGLLIFFGVGCVAALKVAGASFGQWEISVIWGLGVAMAIYLTA  60

E.adnata      HISGAHINPAVTLAFWTSHVFPANKILPYILGQFSGGILASLLL---------------K  105
               +SGAH+NPAVT+A W    F   K++P+I+ Q +G   A+ L+
E.coli        GVSGAHLNPAVTIALWLFACFDKRKVIPFIVSQVAGAFCAAALVYGLYYNLFFDFEQTHH  120

E.adnata      IILGNVANM------GTILPSNDNWLQALIVEIILTFILMFVVL-----GSGIDRRASAS  154
              I+ G+V ++       T    + N++QA  VE+++T ILM ++L     G+G+ R   A
E.coli        IVRGSVESVDLAGTFSTYPNPHINFVQAFAVEMVITAILMGLILALTDDGNGVPRGPLAP  180

E.adnata      FGAIAVGLTVVAEAAFMGPITGAGMNPVRSLAPALVAHIS-------------QYQWLYV  201
                 + +GL +    A MGP+TG  MNP R   P + A ++              Y  + +
E.coli        ---LLIGLLIAVIGASMGPLTGFAMNPARDFGPKVFAWLAGWGNVAFTGGRDIPYFLVPL  237
```

**Fig. S7.** Sequence comparison between *E. adnata* diazoplast and *E. coli* GlcF. Residues in blue and underlines are highly conserved in GlpF available sequences. Residues in red interact with glycerol in the *E. coli* GlcF.

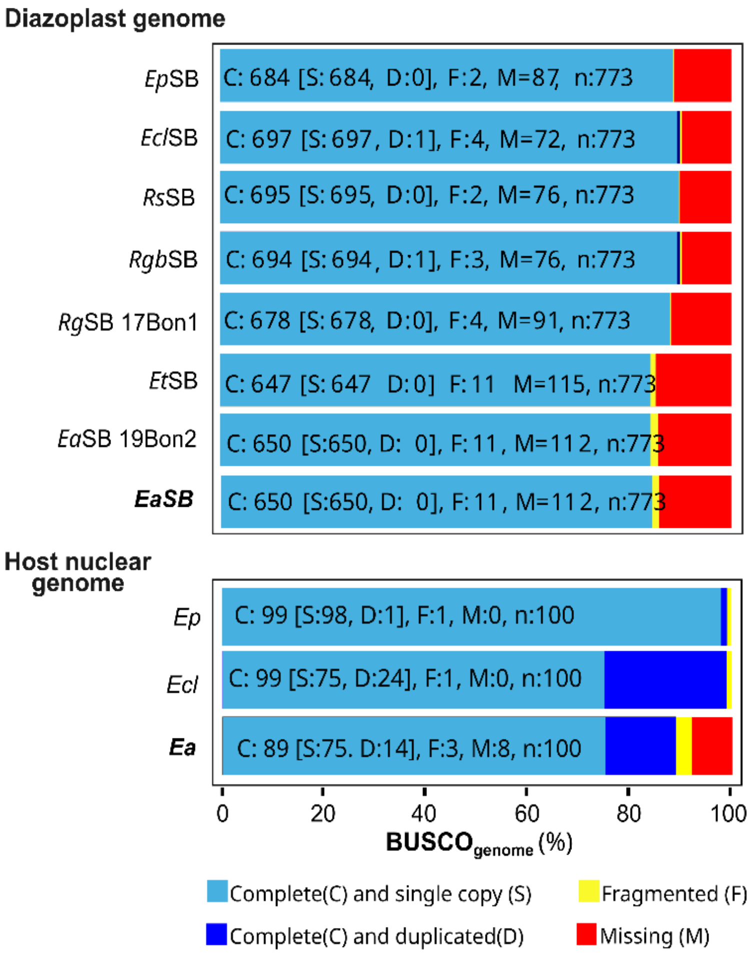


**Fig. S8.** BUSCO (Benchmarking Universal Single-Copy Orthologs) analysis of known diazoplast of nuclear diatom genomes.Diazoplast genome sequence accession: *E. pelagica* (EpSB: NZ_OX373462.1), *E. clementina* (EclSB: CP123991.1), *R. sterrenburgii* (RsSB: CP192514.1), *R. gibberula* (RgbSB: NZ_AP018341.1 ), *R. gibba* isolate 17Bon1 (RgSB 17Bon1: CP067995.1), *E. turgida* (EtSB: AP012549.1), *E. adnata* isolate 19Bon2 (EaSB 19Bon2: CP076462.1), *E. adnata* isolate LB (EaSB, this study). Host nuclear genome sequence accession: *E. pelagica* (Ep; GCA_946965045.2), *E. clementina*; (Ecl; GCA_051400955.1), *E. adnata* isolate LB21(Ea; unpublished).

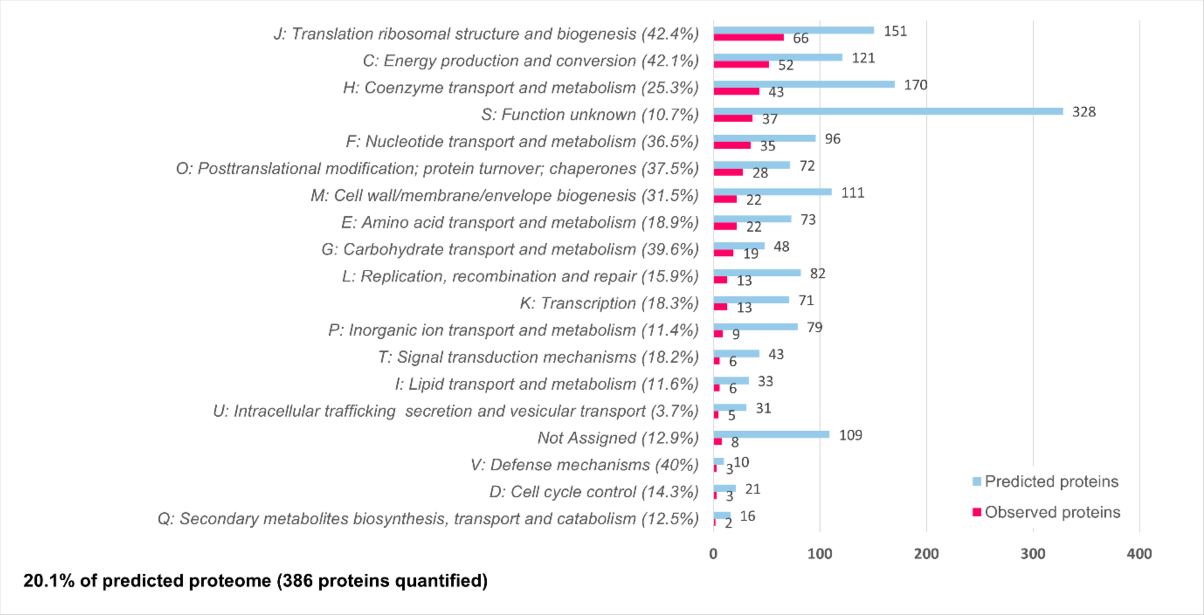


**Fig. S9.** COG categories assigned to predicted and identified *E. adnata* diazoplast proteins

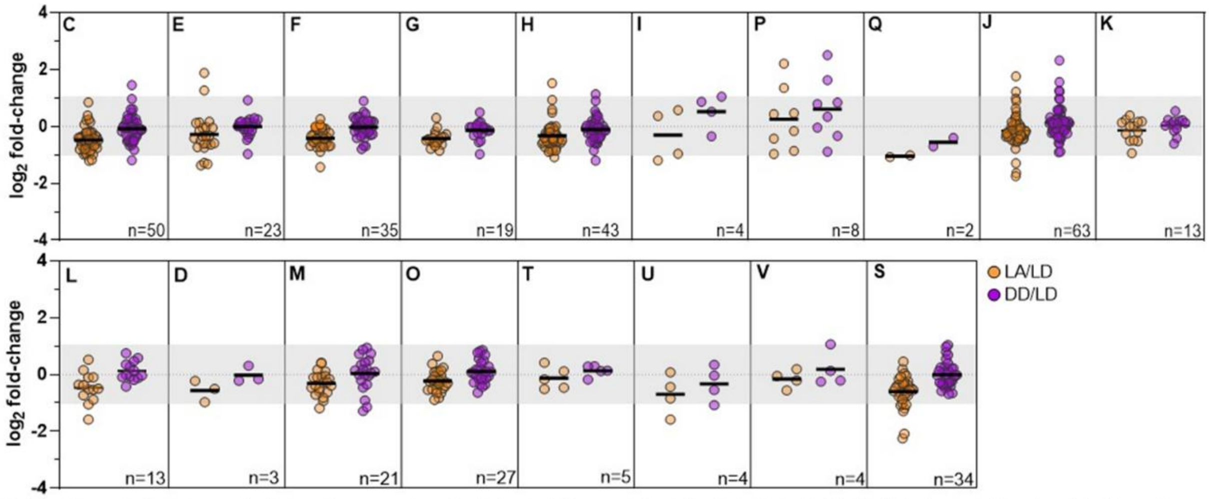


**Metabolism:** **C**, Energy production and conversion; **E**, Amino acid transport and metabolism; **F**, Nucleotide transport and metabolism; **G**, Carbohydrate transport and metabolism; **H**, Coenzyme transport and metabolism; **I**, Lipid transport and metabolism; **P**, Inorganic ion transport and metabolism; **Q**, Secondary metabolites biosynthesis, transport and catabolism.**Information storage and processing:** **J**, Translation, ribosomal structure and biogenesis; **K**, Transcription; **L**, Replication, recombination and repair. **Cellular processes and signaling:** **D**, Cell cycle control; **M**, Cell wall/membrane/envelope biogenesis; **O**, Posttranslational modification, protein turnover, chaperones; **T**, Signal transduction mechanisms; **U**, Intracellular trafficking, secretion and vesicular transport; **V**, Defense mechanisms. **Poorly characterized:** **S**, Unknown function.

**Fig. S10.** Proteome dynamics of the *E. adnata* diazoplast in response to light and $NH_4^+$ supplementation, categorized by COG functional groups. Keys: LD, light-diazotrophic; LA, light-$NH_4^+$; DD, dark-diazotrophic.

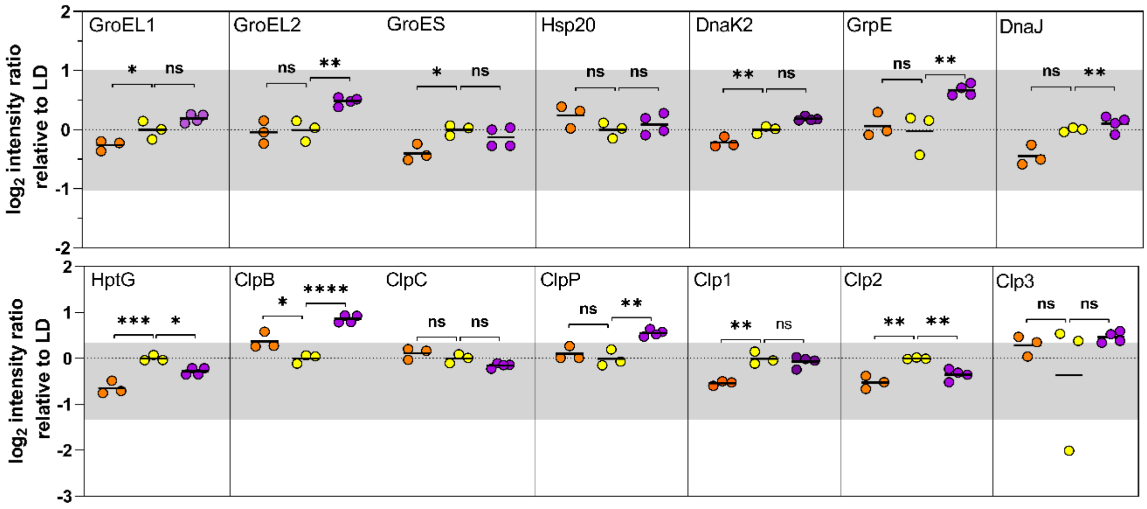


**Fig. S11.** Proteome dynamics of *E. adnata* diazoplast molecular chaperones and proteases under light and $NH_4^+$ supplementation. Keys: LD (light-diazotrophic, yellow), LA (light-$NH_4^+$, orange), DD (dark-diazotrophic, purple). Abundance values were compared to LD, with statistical differences assessed by one-way ANOVA and Dunnett's post hoc test.

```
COP23    aagcgcgtaggttaaggtacgacgactgacgacctgtaaggcg-taatgt(n)84 ATG
Hyp      tatagatgtattattatctactgtaattattaatattcgaacct-taagat(n)295GTG
PetH     caactccgtagacaaaccaactgtttgtctacggagttgtggagttaaatt(n)237ATG
                                  22-23 ntd
```

**Fig. S12.** Putative NtcA-binding sites in *E. adnata* diazoplastic genes responsive to $NH_4^+$ supplementation in the culture medium. The sequence $GTA(N)_8TAC(N)_{23}TA(N)_3T$ represents the consensus binding site of Nitrogen control transcription factor A (NtcA)-activated Class II promoters. Pink boxes highlight the conserved nucleotides in putative promoters of *E. adnata* diazoplastic genes responsive to $NH_4^+$.

**Tabla S1.** List of *E. adnata* diazoplastic proteins responsive to $NH_4^+$ supplementation in the culture medium.

| Protein name | $log_2$ FC | p value | Regulation type |
|---|---|---|---|
| Circadian oscillating protein COP23 | -2.25 | 0.0261 | down |
| Putative competence-damage inducible protein | -2.09 | 0.0014 | down |
| 50S ribosomal protein L29 | -1.75 | 0.0348 | down |
| Phenylalanine-tRNA ligase beta subunit | -1.64 | 0.0341 | down |
| Glucokinase | -1.43 | 0.0047 | down |
| Tryptophan synthase beta chain | -1.32 | 0.0128 | down |
| Xaa-Pro aminopeptidase | -1.28 | 0.0061 | down |
| Cytochrome f | -1.21 | 0.0385 | down |
| 3-oxoacyl-[acyl-carrier-protein] synthase 2 | -1.20 | 0.0007 | down |
| Hypothetical protein | -1.19 | 0.0008 | down |
| Belongs to the peptidase S41A family | -1.18 | 0.0010 | down |
| Plastocyanin | -1.18 | 0.0095 | down |
| O-acetylserine sulfhydrylase | -1.08 | 0.0396 | down |
| PFAM RNA recognition motif | -1.08 | 0.0123 | down |
| Belongs to the UPF0367 family | -1.06 | 0.0067 | down |
| Single-stranded DNA-binding protein | -1.05 | 0.0082 | down |
| Granule associated protein | -1.03 | 0.0035 | down |
| Carboxymethylenebutenolidase | -1.02 | 0.0023 | down |
